# Molecular Simulations and NMR Reveal How Lipid Fluctuations Affect Membrane Mechanics

**DOI:** 10.1101/2022.09.03.506496

**Authors:** Milka Doktorova, George Khelashvili, Rana Ashkar, Michael F. Brown

## Abstract

Lipid bilayers form the main matrix of functional cell membranes, and their dynamics underlie a host of physical and biological processes. Here we show that elastic membrane properties and collective molecular dynamics are related by the mean-square amplitudes (order parameters) and relaxation rates (correlation times) of lipid acyl chain motions. We performed all-atom molecular dynamics simulations of liquid-crystalline bilayers to further interpret available NMR data. Our analysis entailed development of a theoretical framework that allows direct comparison of carbon-hydrogen (CH) bond relaxations as measured by simulations and NMR experiments. The new formalism enables validation of lipid force fields against NMR data by including a fixed bilayer normal (director axis) and restricted anisotropic motion of the CH bonds described by their segmental order parameters. The simulated spectral density of thermally excited CH bond fluctuations exhibited well-defined spin-lattice (Zeeman) relaxations analogous to those measured by solid-state NMR spectroscopy. Their frequency signature could be fit to a simple power-law function, indicative of collective dynamics of a nematic-like nature. The calculated spin-lattice relaxation rates scaled as the squared order parameters of the lipid acyl chains yielding an apparent bending modulus *κ*_C_ of the bilayer. Our results show a strong correlation with *κ*_C_ values obtained from solid-state NMR studies of 1,2-dimyristoyl-*sn*-glycero-3-phosphocholine (DMPC) bilayers with varying amounts of cholesterol as further validated by neutron spin-echo measurements of membrane elasticity. The simulations uncover a critical role of interleaflet coupling in membrane mechanics and thus provide insights into the molecular sites of emerging elastic properties within lipid bilayers.

**STATEMENT OF SIGNIFICANCE:** The lipid make-up of a bilayer determines its measurable properties but how the motions of individual molecules combine to produce these properties remains unclear. By exploiting the synergy between NMR spectroscopy and molecular dynamics (MD) simulations, we show that the lipid dynamics in a bilayer are collective yet segmental in nature and contribute directly to bilayer elasticity. Comparison between MD simulations and NMR entails an improved theoretical framework that allows the two techniques to be directly related. This study thus provides novel insights into the inner workings of lipid membranes while delivering a new tool for validating computational approaches against experimental data.

## INTRODUCTION

Lipid membranes are ubiquitous in biology—they exhibit a large spectrum of physical properties which allows them to serve a multitude of functions (1,2). As the subjects of both experimental and theoretical investigations, they are justifiably of great contemporary interest (2-7). Their rich behavior is enabled by the large diversity of lipid molecules and their distinct mixing properties (8-11). The resulting structural features of the membranes such as bilayer thickness and lipid packing have been studied extensively and are known to influence membrane permeability, elasticity, and protein-membrane interactions (12-17). Lipid molecules also undergo constant thermal motions in the form of orientational fluctuations and segmental dynamics (18). Solid-state nuclear magnetic resonance (NMR) is one of few experimental techniques that can detect and quantify these thermal motions at the level of individual lipid molecules (18-23). It is sensitive to the fluctuations of the lipid acyl chains, in particular the carbon-hydrogen CH (or deuterium) bonds. By measuring the frequency dependence of bond reorientations, i.e., their power spectrum or spectral density, analysis of the bilayer fluctuations is possible through the combination of NMR lineshape and relaxation time measurements. The NMR lineshape directly reveals the various degrees of motional averaging of the molecules and quantifies the order parameters of the carbons along the lipid chains (24-26). At the same time, measurements of the rate of nuclear spin relaxation of each carbon provide information on correlated CH bond fluctuations over multiple time scales (27).

Currently an extensive body of NMR data points to collective dynamics of the lipid molecules that are linked to bilayer mechanical properties (18). However, these results have been challenged by theoretical arguments stemming from the convoluted nature of the spectral density obtained from NMR relaxation measurements. One approach that has the potential to resolve this conundrum is molecular dynamics (MD) simulations. Indeed, acyl chain order parameter profiles obtained from NMR have been historically used to validate the simulation force field parameters (7,28-30). These parameters govern the forces exerted on the atoms in an MD simulation and drive their dynamics, rendering comparisons between theory and experiments particularly informative. While NMR-derived order parameter profiles describe the time-averaged structural properties of the lipid molecules (31), the corresponding relaxation measurements hold the key to understanding the connection between local lipid dynamics and bilayer properties (29,32-34). In particular, the CH bond dynamics show both slow and fast motions, but the origins of this emerging hierarchy of motions remain hidden in the data and pose new questions. Are they a result of the reorientations of individual molecules, as detailed by a so-called molecular model, or rather the concerted motions of larger lipid assemblies—in other words, collective dynamics (18,35)? And if the motions are indeed collective, do they resemble those of nematic liquid crystals where different segments of the lipid chains reorient collectively, or smectic liquid crystals where the identity of the lipids is lost in the dynamical modes of a fluctuating 2D surface? Answers to these questions will not only illuminate the relationship between lipid and bilayer dynamics, but may also inform the correspondence between NMR observables and different experimental techniques that measure global bilayer fluctuations, such as flicker spectroscopy (36). Experimentally validated MD simulations thus present an ideal platform for replicating NMR observables and investigating their molecular origins (29,32,37-39).

Here we use MD simulations to analyze the nature of bilayer dynamics, the relationship between the magnitude of CH bond fluctuations and their relaxation rates, and the resultant elastic modulus as measured with NMR spectroscopy. To facilitate comparison with solid state NMR data, we first develop a theoretical framework for the calculation of spin-lattice relaxation rates from the simulations that overcomes shortcomings of existing numerical approaches. The lipid orientational dynamics are considered for in vitro and in silico systems, together with the inherent anisotropic motion of the CH bond fluctuations. For validation, we simulated a data set of experimentally well-characterized model membranes composed of 1,2-dimyristoyl-*sn*-glycero-3-phosphocholine (DMPC) and cholesterol (40-42). Using NMR-based protocols, we calculate the spectral densities of the lipid CH bond fluctuations from the simulation trajectories and show that they can be modelled by a simple power-law function. Lastly, we evaluate the simulation-derived carbon-specific order parameters and relaxation rates which we find to follow a square-law dependence related to bilayer elasticity. Our findings validate and further clarify interpretations of NMR data while presenting a novel approach for calculating bending moduli of simulated bilayers. Importantly, the simulations identify a previously overlooked region of the bilayer where the two leaflets interdigitate that holds a key to membrane mechanics. Our methodology thus opens new avenues for the validation of lipid force fields in terms of underlying theoretical concepts of membrane elasticity for liquid-crystalline bilayers (5,38,43,44).

## METHODS

### Bilayer construction

Bilayers containing 1,2-dimyristoyl-*sn*-glycero-3-phosphocholine (DMPC) with 0, 20, 33, and 50 mol% cholesterol (Chol) were constructed with the CHARMM-GUI web server (45-49). Each bilayer contained 100 lipids per leaflet and was hydrated with 45 water molecules per lipid with no added salt. The systems thus ranged from 46,000 to 50,000 atoms.

### Simulation protocol

Each system was initially relaxed with CHARMM-GUI’s 6-step equilibration protocol using NAMD version 2.12 (50) and the CHARMM36 force field for lipids (29,51). The simulation temperature was set to 317 K (44 °C), the same value as used in the cognate experimental studies (34,41,52). This initial equilibration was followed by 1 ns of simulation following the CHARMM-GUI simulation production protocol employing a 2-fs time step, 10–12-Å Lennard Jones potential with the NAMD *vdwForceSwitching* option enabled and *rigidbonds* set to all. The particle mesh Ewald (PME) method was used for treatment of the electrostatic interactions with a grid spacing of 1 Å, and a constant temperature (317 K) and pressure (1 atm) maintained with a Langevin thermostat and a Nose-Hoover Langevin piston, respectively. The systems were then transferred to the Summit computer cluster at Oak Ridge National Laboratory, where they were simulated for a total of 2 µs each with the OpenMM software (53). These production simulations implemented PME for electrostatic interactions and were performed at a temperature of 317 K under an NPT ensemble with semi-isotropic pressure coupling. The simulation runs utilized a 2-fs time step and a *friction* parameter set to 1.0/picosecond. Additional parameters included: *EwaldErrorTolerance* 0.0005, *rigidwater* True, *constraints* HBonds, and *ConstraintTolerance* 0.000001. The van der Waals interactions were calculated applying a cutoff distance of 12 Å and switching the potential from 10 Å.

### Simulation analysis

All post-processing analysis was performed with VMD (54), MATLAB (55), and in-house tcl and MATLAB scripts. Unless otherwise noted, calculations were done on all trajectory frames totaling more than 50,000 with output frequency of 40 ps. Each bilayer system was first centered so that in every frame the average position of the terminal methyl carbons of all lipid acyl tails (carbons C14 on the *sn*-1 and *sn*-2 chains, Fig. 1 *A*) was set at (*x,y,z*) = (0,0,0). The average area per lipid ⟨*A*_L_⟩ was calculated by dividing the average lateral area of the simulation box by 100, i.e., the number of lipids in one leaflet, including cholesterol. The acyl chain order parameter (*S*_CD_) at each carbon, defined as the second-rank Wigner rotation matrix element 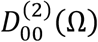 where Ω = (α, β, γ) are the Euler angles (56), was obtained with the formula:

**FIGURE 1:**
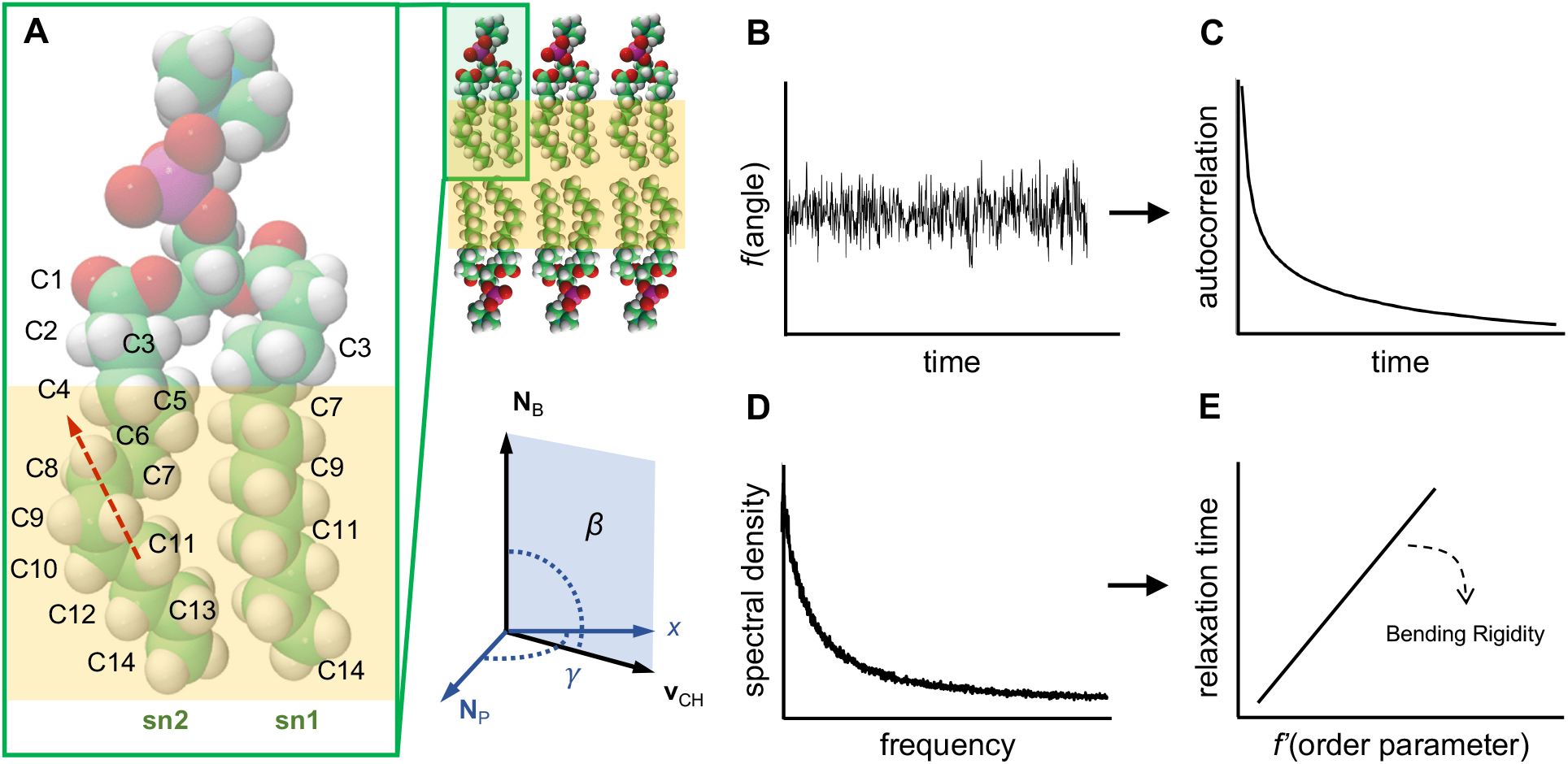
Schematic representation of the simulation methodology. (A) Lipid bilayer and structure of 1,2-dimyristoyl-*sn*-glycero-3-phosphocholine (DMPC). Labels indicate carbons C3, C7, C9, C11, and C14 on the *sn*-1 chain and all carbons on the *sn*-2 chain of the lipid. Carbon atoms are shown in green, hydrogen in white, oxygen in red, and phosphorus in purple. Red dashed arrow shows a local director vector spanning carbons C11 and C8 on the *sn*-2 chain. Yellow shading outlines the part of the lipid chains (in the absence of Chol) where the fluctuations of the CH bonds follow a square-law dependence. Shown also is an example of the angle *β* between a CH bond vector **v**_CH_ on the lipid chain and the bilayer normal **N**_B_ (director axis), as well as the angle *γ* describing rotation of **v**_CH_ around **N**_B_. (B) Fluctuations of *β* and *γ* over time are analyzed to obtain (C) their autocorrelation function averaged across all lipids and time. Fourier transformation of the autocorrelation function gives (D) the spectral density from which (E) the relaxation rate is obtained. Multiple carbon atoms on the lipid chains are used together with their respective order parameters to calculate the bilayer bending modulus. See text for details. **[2 column figure]**

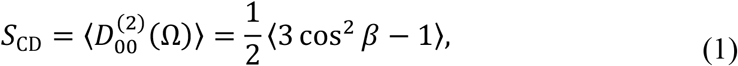

where *β* ≡ *β*(*t*) is the time-dependent angle between a CH bond at this carbon and the bilayer normal (i.e., the *z*-dimension of the simulation box). Likewise, Ω ≡ Ω(t) represents the time-dependent Euler angles and the angular brackets ⟨…⟩ denote an ensemble average over all lipids in the bilayer and all trajectory frames.

### Calculation of autocorrelation functions from simulation trajectories

Autocorrelation functions of the lipid CH bond fluctuations were calculated from the trajectories as follows (Fig. 1 *A–C*). First, the direction of each 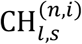 vector in the bilayer was computed as a function of time, where 1 ≤ *l* ≤ *N*_L_ denotes an individual DMPC lipid, 2 ≤ *n* ≤ 14 is the carbon number on the *sn*-1 (*s* = 1) or *sn*-2 (*s* = 2) chain, and *i* = (1,2) is the hydrogen atom. These data were used to calculate three individual time series for each 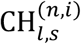 vector **v**_CH_ = (*v*_2_, *v*_3_, *v*_4_), corresponding to the second-rank Wigner rotation matrix elements 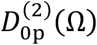 with *p* ∈ [0,1,2], as defined in Ref. (56):

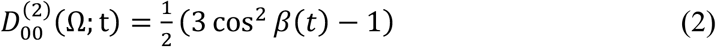

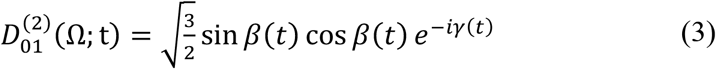

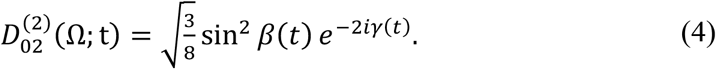

In Eqs. 2–4, 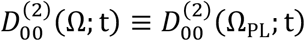 is a function of the Euler angles Ω_PL_ = (α_PL_, β_PL_, γ_PL_) describing the transformation from the principal axis system (PAS) to the laboratory frame (LAB) (see Fig. S1). From this point on, the subscript PL will be suppressed but implied for brevity, unless otherwise noted. Furthermore, *β*(*t*) is the angle between **v**_CH_ and the bilayer normal **N**_B_ = (0,0, ±1) (+1 for the lipids in the top leaflet and −1 for the lipids in the bottom leaflet) at time *t*, and *γ*(*t*) is the time-dependent angle describing the rotation of the 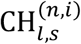 bond with respect to the bilayer normal (see Fig. 1 *A*). Thus, *γ*(*t*) was calculated as the angle between the normal to the plane defined by the cross product of **v**_CH_ and **N**_B_, that is **N**_p_ = (**v**_CH_ × **N**_B_), and the *x*-axis of the simulation box **N**_*x*_ = (1,0,0):

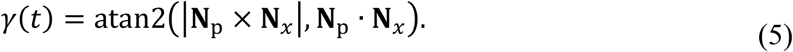

The autocorrelation function of each time series was then obtained as:

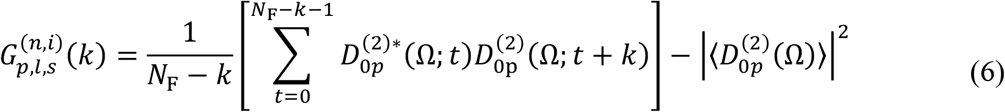

where *N*_F_ is the total number of frames and 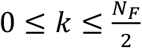 is the lag time (see also Eq. 13 below). Note that the second term on the right in Eq. 6, the square of the mean, vanishes for *p* = [1,2] and is thus nonzero only for 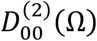. Furthermore, for the special case of lag time *k* = 0, Eq. 6 becomes the variance of 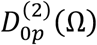:

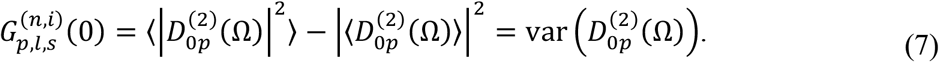

For every carbon 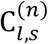 the autocorrelation function from Eq. 6 was averaged over its two covalently bound hydrogen atoms and all DMPC lipids in the bilayer to yield the autocorrelation function for that carbon atom:

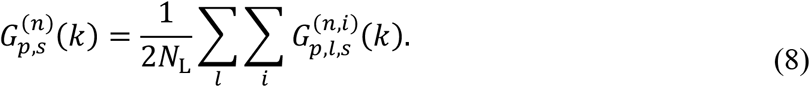

### Theoretical framework for calculation of NMR relaxation rates from molecular dynamics trajectories

The lipid bilayer in the MD simulations corresponds to a membrane patch whose director axis can be assumed to be the average lamellar normal, that is a vector parallel to the *z*-dimension of the simulation box. Hence, the spectral densities of the CH bond fluctuations within this director frame need to first be transformed into the so-called *laboratory reference* system to correspond to the experimental NMR relaxation rates, which are measured in the presence of a magnetic field (see SM and Fig. 4 in Ref. (57)). The director-frame spectral density functions, 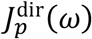, are Fourier transforms of the correlation functions that characterize the internal motions within the membrane. In an NMR experiment, typically one can assume that the relaxation rates are orientationally averaged, because for the case of lipid multilamellar dispersions rapid orientational averaging occurs over all the director orientations during the relaxation (∼50–100 ms and longer). Furthermore, if one assumes unrestricted *isotropic* motion with a single correlation time (the solution NMR limit), the relaxation rate *R*_1Z_ simplifies to:

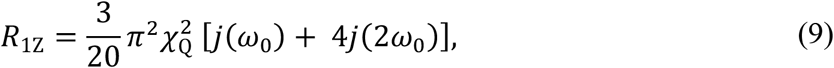

where 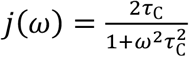 is a Lorentzian reduced spectral density with *τ*_C_ as the correlation time, *χ*_Q_ is the static quadrupolar coupling constant, and *ω*_0_ = 2*πv*_0_ is the Larmor frequency of the NMR measurement (24,35) (see Eq. S22 and Discussion).

However, in a bilayer simulation run with periodic boundary conditions and semi-isotropic pressure coupling, the lamellar normal (director) remains fixed and the assumption for unrestricted isotropic motion and thus Eq. 9, does not hold. Instead, to compare the simulation results to the experimental ones, the orientationally averaged but *anisotropic* spin-lattice relaxation rate needs to be expressed in terms of the director-frame spectral densities (see Discussion and SM for details of the derivation):

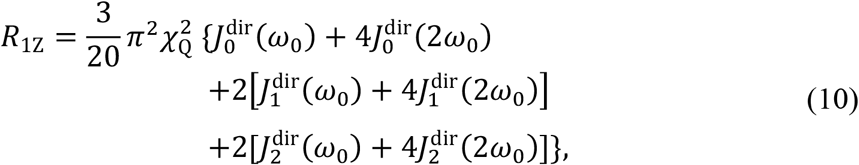

where:

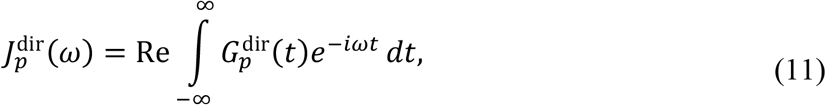

and:

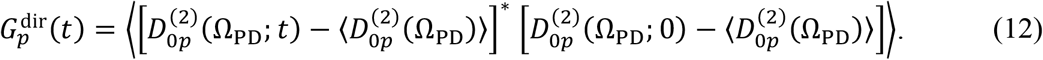

The value of the numerical pre-factor in Eq. 10 is 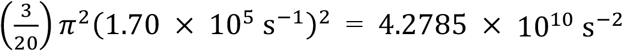 (see SM) and the spectral density 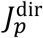 is defined as the two-sided Fourier transform of the correlation function 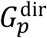. The latter decays to a zero value because the fluctuations are expressed relative to the average values of the Winger rotation matrix elements. The director-frame correlation functions can also be written by subtracting the modulus-squared of the average value to read:

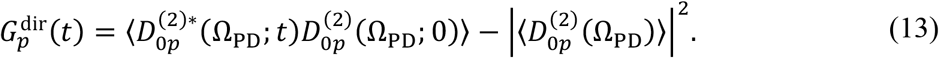

Note that for a cylindrically symmetric distribution, the last term on the right becomes 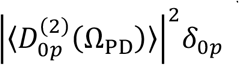 where 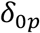 is the Kronecker delta function. For completeness, the orientationally averaged relaxation rates can also be expressed in terms of the spherical harmonic correlation functions. The equivalent results in terms of the spherical harmonic correlation functions and spherical harmonic spectral densities can be found in the SM.

### Calculation of relaxation rates from the simulation trajectories

Equations 10–13 provide the basis for the calculation of the relaxation rates from the simulation trajectories. Note that the autocorrelation function 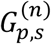 for the molecular dynamics simulation trajectories in Eq. 8 is the same as the correlation function 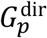 in Eq. 13. We can thus take its discrete Fourier transform to arrive at an expression for the spectral density 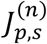 of the 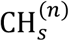 bond fluctuations from the simulations:

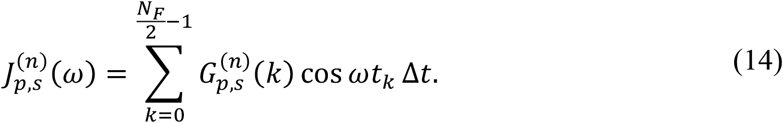

In Eq. 14 *ω* = 2*πv* with *v* being the Larmor frequency, Δ*t* is the sampling time interval, and *t*_*k*_ = *k*Δ*t* is the time at lag *k*.

Equation 14 is a one-sided Fourier transform while the continuous Fourier integral in Eq. 11 goes from [−∞, +∞]. We therefore need to multiply the former by a factor of two but without overcounting the element at *k* = 0, which thus reads:

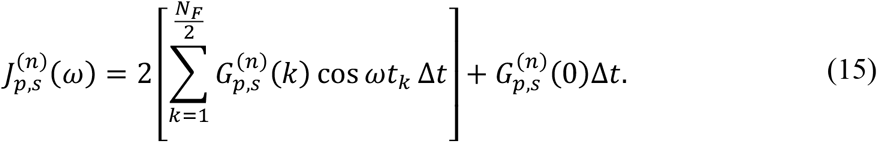

As mentioned above, the *t* = 0 element of the correlation function 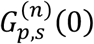 is the variance of 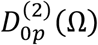 Eq. 7). Accordingly, the second term in Eq. 15 represents a constant equal to 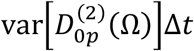 which is added to 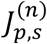 at every frequency. Estimation of the spectral density from the simulations thus exhibits a strong dependence on the discrete time step Δ*t* (sampling interval). Normally, Δ*t* could be as small as the frequency with which atomic coordinates are written out during the simulation if every single frame is used for analysis (in our case Δ*t* = 40 ps); alternatively, if every *n*th frame is used instead, then the sampling frequency would be *n*Δ*t*. Note that the simulations are run with a 2-fs timestep which puts a lower bound on Δ*t*, but such frequent output is in general impractical for microsecond-long simulations.

Having this Δ*t*-dependent constant would result in an unrealistic shift of the spectral density and consequently, the relaxation rates. Thus, to remove this apparent dependence on Δ*t* and recover the analytical result in Eq. 11, which is in the limit of Δ*t* → 0 (see Discussion), we first fit 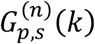 to a well-defined function, and then use the fit to resample the correlation function at a much smaller Δ*t*_fit_ ≪ Δ*t*. For all CH bonds the time series of the second-rank Wigner rotation matrix elements from Eqs. 2–4 follow a power law of the form: *ax*^*b*^ + *c* (see Fig. 2 *A* and Figs. S1–S3). We therefore fit 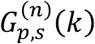 for *k* ≥ 1 to a power-law function and use the best fit 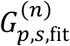 together with the variance 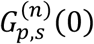 at *k* = 0 to resample the correlation function at a smaller Δ*t*_fit_ interval and recover the spectral density:

**FIGURE 2:**
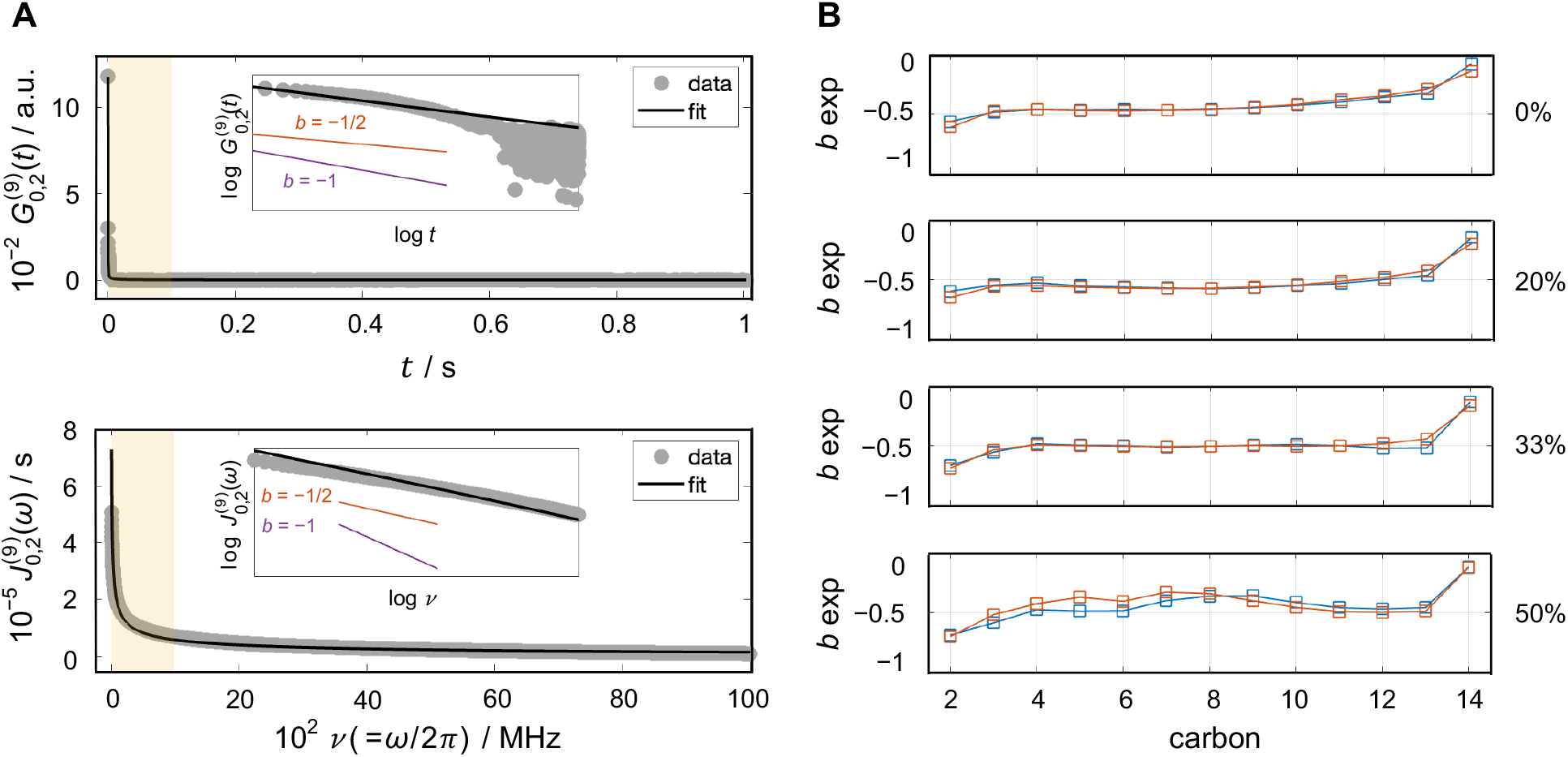
Dynamics of CH bond fluctuations. (A) Example of (top) autocorrelation function 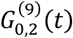 and (bottom) spectral density function 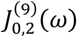 for carbon C9 of the *sn*-2 chain of DMPC. The results correspond to fluctuations of the CH bonds as described by the Wigner 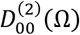 function. Simulation data are shown in grey and best power-law fit to the simulation results in black. Insets show expansion of the data from the highlighted region from 0–1000 MHz but in a log-log plot. Shown for comparison are sample functions of the form *ax*^−1^ (purple) and *ax*^−1*/*2^ (red). (B) Power exponents of best fits to the spectral densities 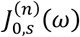 carbons C2–C14 on the *sn*-1 (red) and *sn*-2 (blue) chains of DMPC with 0, 20, 33, and 50 mol % cholesterol (Chol). The power exponents of all carbons in all simulated bilayers are close to −1/2 consistent with collective segmental dynamics. All simulations were performed at 44 °C. See text for details. **[2 column figure]**

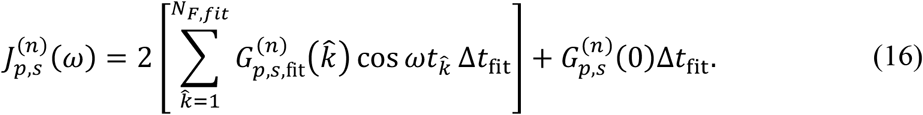

Note that Eq. 16 is calculated at 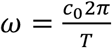 (or equivalently, 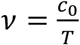) where 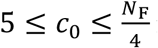 is an integer and 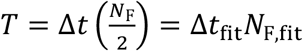 is the total sampled time in accord with the Nyquist-Shannon sampling theorem. The choice of Δ*t*_fit_ is constrained by the variance of 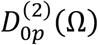 which is the first and largest element of 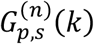. Therefore, to ensure the smoothness of the reconstructed correlation function, for each carbon 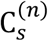 we set Δ*t*_fit_ equal to the minimum value Δ*t*′ such that Δ*t*_fit_ = min (Δ*t*′) where 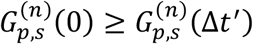. Figures S4–S5 show the reconstructed correlation functions and resulting carbon Δ*t*_fit_ values (see Discussion).

Accordingly, the theoretical spectral densities 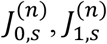, and 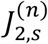 from Eq. 16 are fit to power-law functions of the form *ax*^*b*^ + *c* and the best fits are then used to calculate the relaxation rate of carbon 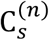 following Eq. 10:

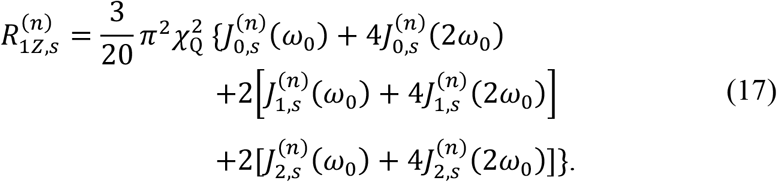

In the above formula *χ*_*Q*_ = 170 kHz, *ω*_0_ is the resonance (Larmor) frequency, and all other symbols are previously defined. The theoretical spectral densities 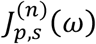 refer to the CH fluctuations with respect to the bilayer normal (director axis) in terms of their mean-square amplitudes (variance) and reduced values. The theoretical relaxation rates are averaged over all director orientations to correspond to the actual experimental measurements. Importantly, although the relaxation rates are orientationally averaged, they are not the same as the isotropic solution NMR results found in textbooks. Use of the latter (obtained by application of the spherical harmonic addition theorem) is often assumed but is inapplicable to lipid membranes because of the underlying assumption that the fluctuations are isotropic, i.e., there is no order parameter. Rather, the CH bond fluctuations must be expressed with respect to the bilayer director and occur with restricted amplitude, as described by their orientational order parameters. Thus, the formula applicable to MD simulations and to the validation of MD force fields based on NMR relaxation is given in Eq. 17 above. This result is based on group theory principles (35) and is different from what is obtained by application of the spherical harmonic addition theorem (see Discussion). Validation of the theoretical and experimental results is further discussed below. Note that the mean-square amplitudes are equal to the area under the spectral density in the region [0, ∞] and thus cannot be numerically determined from the integral of 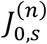 due to the finite sampling. Instead the mean-square amplitudes are thought of as the orientational order parameters in terms of a Clebsch-Gordon series expansion (35), and can be calculated from the raw simulation data (i.e., the time series of the Wigner rotation matrix elements from Eq. 2–4).

### Application of NMR-based approach to extract apparent bilayer bending rigidity from MD trajectories

An apparent bending modulus *κ*_C_ can be extracted from the slope of the theoretically predicted square-law dependence between the relaxation rate *R*_1Z_ and order parameter *S*_CD_ of the acyl chain carbons, as derived previously (35). Following the same NMR-based protocol, we obtain *κ*_C_ of the simulated bilayers with the formula:

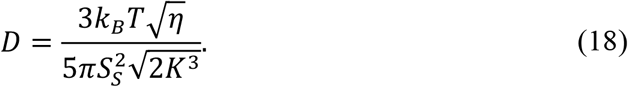

In Eq. 18 *η* = 0.1 Pa · s is the bilayer viscosity, *S*_S_ = 0.6 is the order parameter estimated for slow motions, 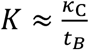 is an elastic constant related to *κ*_C_ and the *full* bilayer thickness *t*_=_ (see below), and *D* is a constant in units of 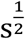 defined as:

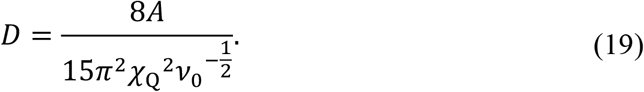

In Eq. 19 the constant *A* is the slope of the dependence of *R*_1Z_ on |*S*_CD_|^2^, *χ*_Q_ is the static quadrupolar coupling constant, and *v*_0_ = 76.8 MHz is the deuterium Larmor frequency. The thickness *t*_=_ which is used to obtain *K* in Eq. 18 represents the full bilayer thickness and is calculated from the number density profile of the bilayer. Specifically, a single number density profile of all bilayer lipid atoms is calculated with the Density Profile plugin in VMD (58) for each system. The profile has two peaks corresponding to the maximum density in the hydrocarbon regions of the two leaflets and decreases to 0 at the ends of the headgroup regions where bulk water begins (see Results). The thickness *t*_=_ used in Eq. 18 is set equal to the distance between the two tails of the bilayer number density profile where the density drops below 5% of the maximum (peak) density. Table S1 lists the corresponding thicknesses for all simulated bilayers.

The full range of carbons used for the square-law fits in the simulations included C4–C13, C6– C13, C8–C13, and C10–C13 for the bilayers with 0, 20, 33, and 50% cholesterol, respectively. Errors for the *κ*_C_ values obtained from the fits were calculated as the standard deviation of the slopes resulting from the best fits to the data after excluding the last 0, 1, and 2 carbon atoms with largest order parameters from each chain. Note that the terminal methyl carbon C14 was excluded from the fits due to its different geometry.

### Splay-fluctuations-based analysis of bilayer bending rigidity

The *κ*_C_ modulus can also be obtained in simulations of lipid bilayers from analysis of the fluctuations in lipid splay angles (see (59) and references therein). Briefly, the procedure involves calculating the probability distribution of splay angles (i.e., the angles between the director vectors) of neighboring pairs of lipids within a leaflet and using this distribution to obtain a potential of mean force (PMF) profile. A quadratic function is then fit to the PMF in the region of small deviations from the mean, and the coefficient of the quadratic term is the corresponding splay modulus for that lipid pair. The splay moduli of all lipid pairs are subsequently weighted and summed, revealing the leaflet *κ*_C_.

The bilayer *κ*_C_ is then the sum of the *κ*_C_ values of the two opposed monolayers. Here, we used the OpenStructure software and algorithm presented in Refs. (60,61) to calculate *κ*_C_ with an alternative computational method from the splay moduli of DMPC, cholesterol (Chol), and DMPC-Chol lipid pairs in the simulation trajectories.

### Analysis of local director fluctuations

Local director (LD) vectors of different lengths were defined at each carbon along the lipid acyl chains. Each LD vector connects the carbon of interest 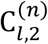 on the *sn*-2 chain either to another carbon atom 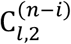 further up the chain on the same lipid *l* towards the lipid headgroup with 1 ≤ *i* ≤ 12, or to the lipid phosphorus atom P (see Results). The fluctuations of the orientations of an LD vector were analyzed in a manner similar to the calculation of the spectral densities described in Eqs. 8 and 15. The only differences are that all angles are defined with respect to the LD vector instead of the CH bond, the denominator in the pre-factor in Eq. 8 is *N*_L_ instead of 2*N*_L_, and since the autocorrelation functions of the LD vector fluctuations do not follow a power law or an exponential function, no fitting and resampling of the correlation was performed. Instead, the spectral density was calculated with Eq. 15 using the sampling interval corresponding to the simulation Δ*t* of 40 ps.

### Measurement of bending moduli with neutron spin echo spectroscopy

To extend the results of the MD simulations and NMR analysis, complementary neutron spin echo (NSE) measurements were performed on unilamellar vesicles extruded through a polycarbonate filter with 100-nm pore diameter (see SM for more details). The NSE data yielded the normalized intermediate scattering function, *I*(*q, t*)*/I*(*q*, 0), for discrete *q*-values within the accessed *q*-range, where *t* is the Fourier time. For lipid membranes, the probed dynamics follow a stretched exponential function with a stretching exponent of 2/3, such that (62):

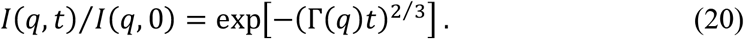

Fits of the intermediate scattering functions using the above equation yield the decay rate, Γ(*q*), at individual *q*-values. Plots of Γ(*q*) versus *q* show the typical *q*^3^ dependence for thermally undulating elastic thin sheets predicted by Zilman and Granek. Using theoretical refinements by Watson and Brown (63), based on the Seifert-Langer model (64), allow calculations of a renormalized bending rigidity 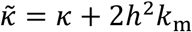, where *h* is the height of the neutral surface from the midplane and *k*_m_ is the monolayer area compressibility modulus. Assuming the neutral plane to be at the interface between the hydrophilic headgroups and the hydrophobic tails, these refinements result in a modified expression of the Zilman-Granek decay rates, Γ(*q*), given by (65):

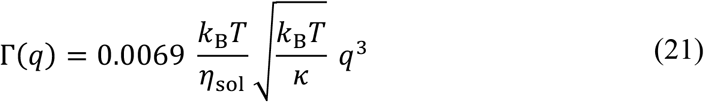

Here, *k*_B_*T* is the thermal energy and *η*_sol_ is the solvent (i.e. D_2_O buffer) viscosity.

## RESULTS

### Lipid dynamics at length scales smaller than the membrane thickness resemble nematic liquid crystals

Membrane lipid molecules are known to diffuse rapidly within the plane of a fluid bilayer while their chains explore various rotational degrees of freedom (29,37,66,67). These thermally excited motions are governed by both bonded and non-bonded interactions and can be rather inhomogeneous along the lengths of the lipid chains due to the hydrophobic effect holding the two leaflets together (68). Nuclear spin relaxation measurements show both fast and slow components of the CH bond fluctuations at each carbon (34). Still, because the resulting signal has contributions from all lipids in the bilayer, the origins of this hierarchy of motions, and consequently the nature of the lipid dynamics needs further investigation (27,32,38,39,69). To help address this question, we sought to examine the thermal fluctuations of the CH bonds along the lipid acyl chains with all-atom MD simulations (59,66,70). As a test system we chose DMPC bilayers with increasing amounts of cholesterol that have been well characterized experimentally (34,41,52). In the past, CH bond fluctuations from simulation trajectories have been expressed in terms of spherical harmonics using spherical polar angles (71,72). However, that representation presents certain challenges (see Discussion) and we sought to derive a theoretical framework centered around the so-called Wigner rotation matrix elements in terms of Euler angles instead (35,73,74). The latter represent functions (Eqs. 2–4) of the Euler angles describing the CH bond orientation: angle *β* between the CH bond and the bilayer normal and angle *γ* which describes the rotation of the CH bond with respect to the bilayer normal (see Fig. 1 and Eq. 5). The autocorrelation function of the time series of each of these functions describes a different relaxation mode of the CH bond fluctuations and was found to follow a power-law dependence (see Fig. 2 *A* and Methods). Its Fourier transform yields the spectral density as a function of frequency that can be directly related to the relaxation rates measured by NMR spectroscopy (Eq. 17).

Importantly, the functional form of the spectral density contains information about the nature of the bilayer dynamics. Each spectral density function calculated from the simulations was found to follow a power-law dependence of the form *y* = *ax*^*b*^ + *c* (Fig. 2 *A* and Figs. S6–S8). This functional form has been shown to correspond to collective dynamics of the lipids, and its power-law exponent represents the dimensionality of the collective lipid interactions formulated in terms of quasi-elastic order-director fluctuations (ODF) (35). At this level, a continuum of wave-like disturbances with an exponent of −1 indicates 2D smectic-like dynamic undulations (20,73,75-77) while an exponent of −1/2 points to 3D nematic-like fluctuations (24,35,78). Figures 2 *B* and S9 show the exponent *b* for all carbons on the *sn*-1 and *sn*-2 chains of the lipids in the simulated DMPC bilayers. As seen in the plots, the power exponents are very close to −1/2 throughout both chains in all trajectories. This result suggests that the dynamics of the lipid hydrocarbon acyl tails are collective, and yet locally more segmental in nature, resembling the behavior of nematic liquid crystals at distances less than the bilayer thickness (79).

To explore this interpretation further, we compared the fluctuations of the lipid CH bonds to those of local director (LD) vectors of varying lengths (Fig. 3). An LD vector connects a carbon either to another carbon at increasing distance further up the chain, or to the phosphate atom (red dashed arrow in Fig. 1 *A* and arrows in the schematic in Fig. 3). The spectral density of the fluctuations of each LD vector again follows a power law (Figs. S10–S12). Fits to a power-law function yielded the corresponding *b-*coefficients. Figure 3 exemplifies the relationship between these *b-*exponents and the length of the LD vectors by showing *b* versus length of the director vectors originating from carbons C7, C9, and C11 on the *sn*-2 chain of DMPC (Fig. 1 *A*). As seen from this figure, an increase in the length of an LD vector results in more smectic-like dynamics as the power exponent decreases, eventually approaching a value of ∼ 0.7. Yet, the motions of the CH bonds resemble most closely those of an LD vector of length 3–4 carbons (compare where the dashed lines cross the corresponding data points in Fig. 3), in agreement with a model of nematic-like dynamics of relaxation rates and order parameters, as originally proposed (35,79).

**FIGURE 3:**
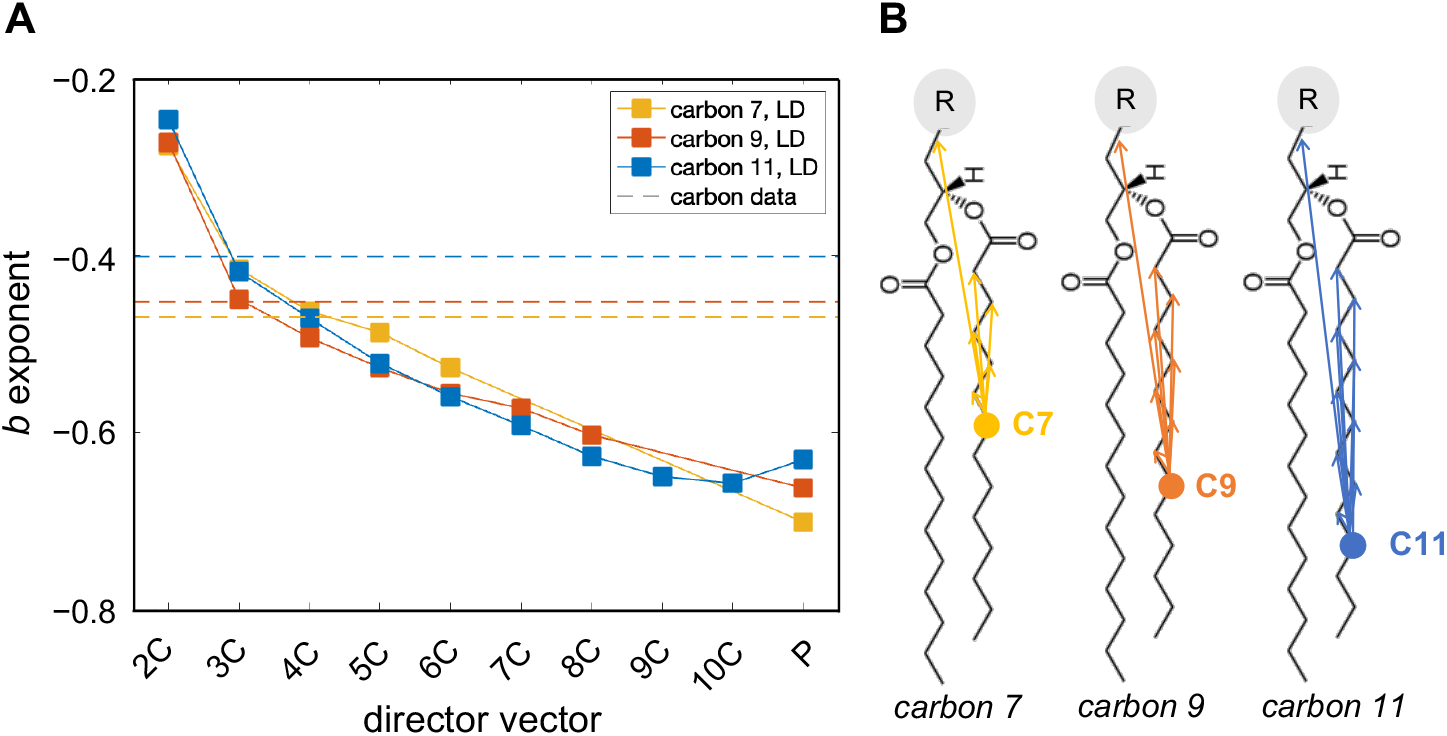
Dynamics of local director (LD) vectors connecting two carbons (or a carbon and the phosphate) of a lipid. (A) Plotted are the *b*-exponents of the best power-law fits to the spectral densities of LD vectors originating from C7 (yellow), C9 (red), and C11 (blue) carbons on the *sn*-2 chain of DMPC in the cholesterol-free bilayer. The LD vectors have lengths between two (2C) and ten (10C) carbons, or extend all the way to the phosphorus atom (P). Shown for comparison are the exponents of the best power-law fits to the spectral densities of the CH bonds at C7, C9 and C11 carbons (dashed lines). (B) Schematic of all analyzed LD vectors. The CH bond fluctuations correspond most closely to LD vectors of length 3–4 carbons, illustrating the segmental nature of the CH bond dynamics. See text for details. **[1.5 column figure]**

### Square-law dependence holds for specific carbon atoms

As mentioned earlier, the spectral densities of the CH bond fluctuations of a carbon atom on the lipid chains can be used to calculate the spin-lattice relaxation time *R*_1Z_ of the bond fluctuations (Eq. 17, Fig. 1). Various NMR experiments have shown that often in bilayers *R*_1Z_ is linearly dependent on the square of the order parameter, a rule thus termed the square-law (35,41,80). To understand the molecular origin of this surprising relationship — and test if it holds in simulated bilayers — we analyzed this trend in our trajectories of DMPC bilayers with increasing amounts of cholesterol (0, 20, 33, and 50 mol %). As expected, the addition of cholesterol made the bilayer more ordered and tightly packed, decreasing the average area per lipid and increasing the bilayer thickness (Fig. S13). From this set of simulations, we calculated the spectral density profiles of the CH bonds along the two chains of DMPC in each trajectory and used them to obtain the corresponding relaxation rates at the NMR deuterium frequency of 76.8 MHz, corresponding to a magnetic field strength of 11.7 T (Eq. 17). Figure 4 shows the relaxation rates *R*_1Z_ (see Methods) and order parameters for all carbons in the simulated bilayers with 0 and 50 mol % cholesterol, and how the two variables are functionally related to each other. From the corresponding log-log plots (Fig. S14) it becomes apparent that they exhibit a square-law relationship for a subset of the carbon atoms in the lower parts of the chains. In particular, carbons C4 through C13 in the DMPC bilayer without cholesterol and carbons within C6 through C13 in the DMPC/Chol bilayers with 20, 33, and 50% cholesterol, respectively, (Fig. 3 *B*, Fig. S14) clearly follow the square-law dependence, consistent with the experimental findings (41). The observed behavior is not chain-dependent, as we see the same dependence for the *sn*-1 and *sn*-2 chains of the lipids (Fig. S14). Note that the C14 terminal methyl carbons on both chains have a different geometry and are therefore excluded from the analysis.

**FIGURE 4:**
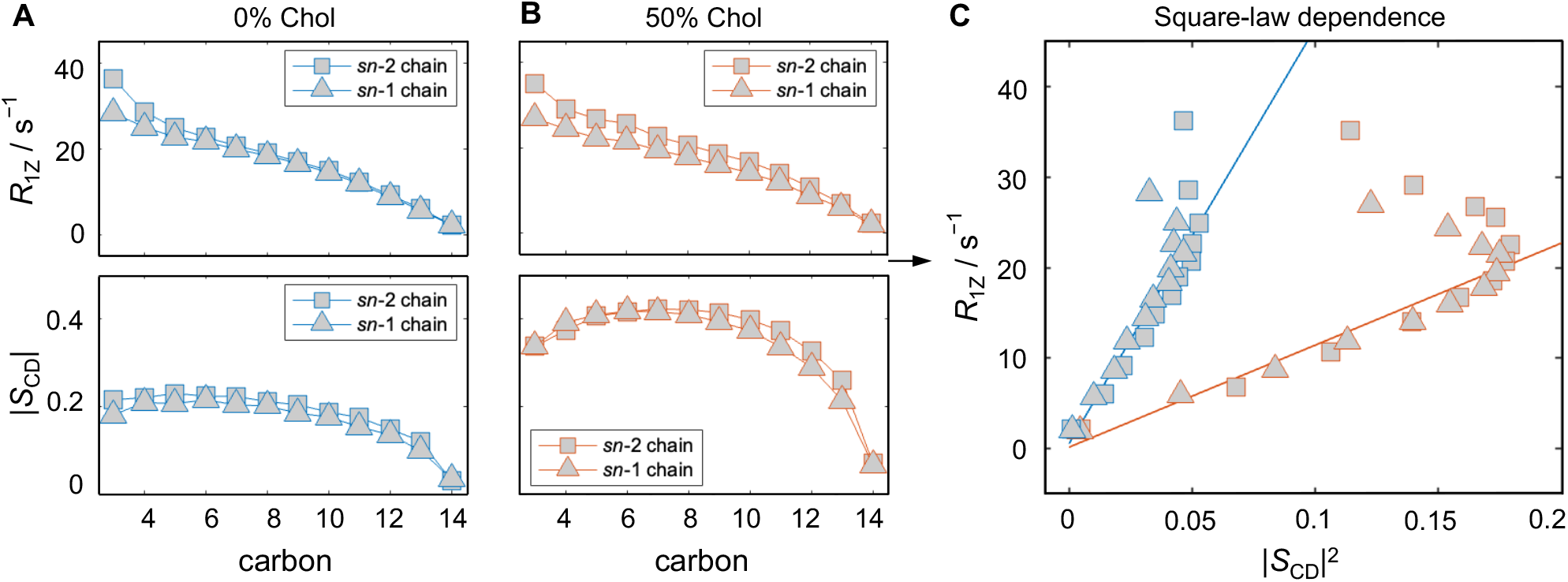
Functional dependence of relaxation rates on CH bond order parameters in simulated bilayers. Data are shown for: (A) DMPC bilayer in the absence (blue) of cholesterol and (B) DMPC containing 50 mol% cholesterol (red). Profiles of spin-lattice relaxation rates (*R*_1Z_, top) and order parameters (*S*_CD_, bottom) are plotted as a function of carbon position along the *sn*-1 and *sn*-2 chains of DMPC. (C) Same data but now re-plotted as the *R*_1Z_ relaxation rate as a function of the squared *S*_CD_ order parameter. Best linear fits to the respective carbon atoms in each bilayer are shown in blue and red, respectively. Note that the relaxation rates and order parameters exhibit a square-law dependence both with and without cholesterol. All simulations were performed at 44 °C. See text for further details. **[2 column figure]**

Interestingly, the square-law in the simulated bilayers breaks down for carbons further up the chain (Fig. 4, Fig. S14). These carbons are schematically illustrated in Fig. 1 *A* and reside at the top part of the acyl groups. To examine in more detail the differences between these carbons and the ones that follow the square-law, we calculated the average position of the carbon atoms relative to the bilayer midplane (Fig. 5 *A*) and compared their number density distributions (Fig. 5 *B*). These data show that regardless of the presence of cholesterol, the carbons that follow the square-law are consistently those in the middle of the bilayer, whose average depth within the membrane falls linearly with their squared order parameters. Stated differently, the linear relationship between *R*_1Z_ and |*S*_CD_|^2^ holds precisely for the CH bonds residing within the region of the bilayer sufficiently far from the headgroups where the two leaflets intercalate (Figs. 1 and 5). Thus, the existence of the square-law dependence uncovers the difference between interfacial (i.e., closer to the water–hydrocarbon interface) and non-interfacial (i.e., closer to the bilayer midplane) dynamics. While both are modulated by collective motions, the former are more smectic-like and the latter are more nematic-like in nature. These observations provide a possible explanation for the mechanistic origin of the square-law relationship and why it holds mainly for a subset of carbon atoms in the lipid acyl chains (see Discussion).

**FIGURE 5:**
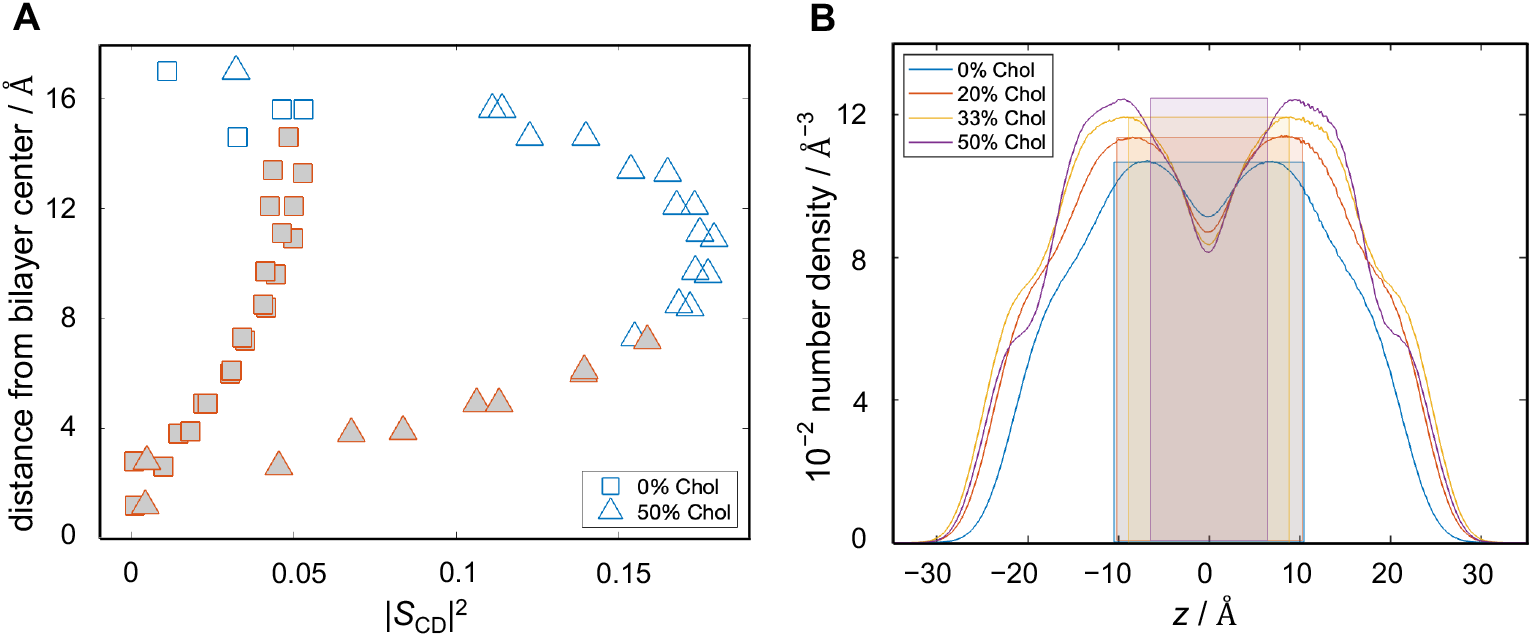
Properties of acyl carbons that follow square-law dependence of relaxation rates and order parameters in simulated bilayers. (A) Distance from the bilayer center as a function of the squared order parameter |*S*_CD_|^2^ for each carbon in the bilayers with 0 and 50% cholesterol. Filled symbols indicate the carbons used for the square-law fits. (B) Number density profiles including all bilayer atoms of the simulated bilayers. Color-coded highlighted areas show the regions of the carbon atoms used in the square-law fits for the corresponding bilayers. The carbons that follow the square-law dependence are within the region where lipid dynamics are influenced by interleaflet interactions. All simulations were performed at 44 °C. **[1.5 column figure]**

### Bilayer bending modulus emerges from the square-law dependence

It has been previously shown that the square-law dependence, i.e., the linear relationship between *R*_1*Z*_ and |*S*_CD_|^2^, is related to the elastic properties of the bilayer in the liquid-crystalline state (35). In particular, the authors used the slope emerging from the fit of the data to estimate an apparent *κ*_C_ bilayer bending modulus. The calculation involved three parameters, namely the bilayer viscosity, the order parameter for slow motions, and a quadrupolar coupling constant, in addition to the bilayer thickness which is proportional to *κ*_C_ (see Eqs. 18, 19). Accordingly, having shown that the square-law holds in our atomistic MD simulations, we followed the same protocol as in Ref. (81) to estimate *κ*_C_ of the simulated bilayers. Figure 6 *A* shows the calculated values for *R*_1*Z*_ and |*S*_CD_|^2^ for DMPC with 0 and 50 mol % cholesterol from the simulations (green), as well as NMR data for the identical systems together with their corresponding best linear fits (black). There is a very good agreement between theory and experiment, confirming the ability of the simulations to recapitulate the properties of the experimental model system.

**FIGURE 6:**
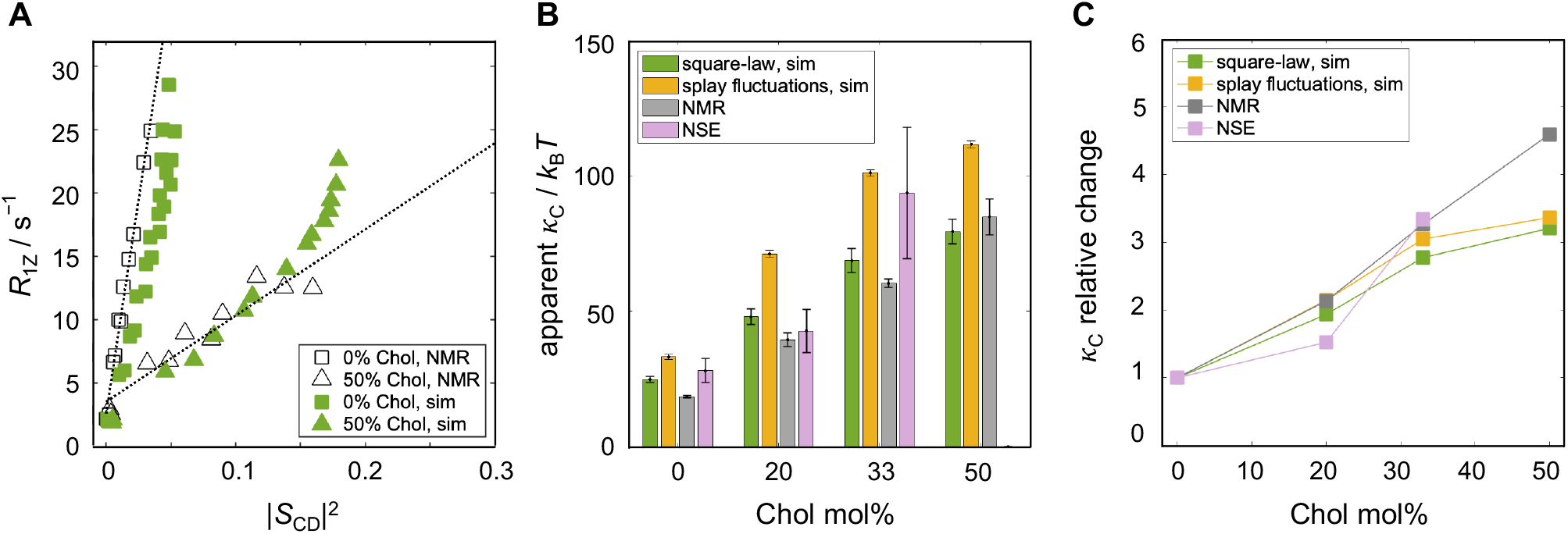
Square-law dependence yields an estimate of bilayer bending rigidity. (A) Simulation (green) and NMR (black) data and best square-law fits to the experimental data for DMPC (squares) and DMPC/Chol 50/50 (triangles) in the liquid-crystalline state. Experimental data was taken from Ref. (41) and simulation data is shown for carbons C4 through C14 for DMPC and C8 through C14 for DMPC/Chol. (B) Comparison between the apparent bilayer bending modulus *κ*_*C*_ calculated from the square-law relationship (green) or splay fluctuations (yellow) in the simulations. Shown also are experimental results for *κ*_*C*_ obtained from the square-law relationship of NMR data (gray) and from bilayer thickness fluctuations measured with NSE (purple). All *κ*_*C*_ values are in units of *k*_B_*T*. (C) Changes in *κ*_*C*_ in the cholesterol-containing bilayers relative to the bilayer without cholesterol. Plotted is the ratio between the two for each of the four methods displayed in panel (B). Errors for the *κ*_*C*_ values obtained from the experimentally determined slope of the square-law relationship were calculated as the standard deviation of the slopes resulting from the best fits to the data after excluding the last 0, 1, and 2 carbon atoms with largest order parameters. The corresponding errors from the square-law dependence in the simulations were calculated in a similar way, see Methods. All simulations were performed at 44 °C. The functional dependence between relaxation rates and order parameters is directly related to the bilayer bending rigidity as validated for the full range of cholesterol concentrations. **[2 column figure]**

Using previously estimated values of 0.1 Pa · s for the bilayer viscosity, ∼0.6 for the slow order parameter, and 167–170 kHz for the quadrupolar coupling constant (56), as well as the full bilayer thickness (see Methods and Table S1), we calculated the corresponding *κ*_C_ values as outlined in the Methods section. The resulting bending moduli are shown in Fig. 6 *B*. As expected, the apparent bilayer *κ*_C_ from the simulations increases with increasing cholesterol concentration, starting at 24.8 (± 1.1) *k*_B_*T* for the pure DMPC membrane and reaching 79.5 (± 4.5) *k*_B_*T* at 50 mol % cholesterol. For comparison, we used an alternative computational method to obtain *κ*_C_ from the simulation trajectories, namely by analyzing the fluctuations in lipid splay angles as described in detail in Refs. (60,61) and summarized in the Methods section. As shown in Fig. 6 *B*, while the absolute values are different, the two computational methods show almost identical trends with cholesterol concentration (Fig. 6 *C*). Plotted in the same figure are also the respective bending rigidities from NMR calculated from the slopes of the experimental data (41) with the same parameters (viscosity, slow order parameter, quadrupolar coupling constant, and thickness) used in the simulations analysis. Not surprisingly, considering the strong agreement of the raw data (Fig. 6 *A*), the NMR results are very close to the simulation ones (Fig. 6 *B*). To obtain an independent experimental validation of the bending moduli, we also used neutron spin-echo (NSE) spectroscopy to analyze the thickness fluctuations of extruded DMPC liposomes with 0, 20, and 33 mol % cholesterol. As shown previously and summarized in Methods, these thickness fluctuations are related to the bilayer bending rigidity via the Zilman-Granek theory (62). The results show a very good overall agreement with the computational methods, with a clear stiffening effect of cholesterol (Fig. 6 *B, C*), confirming the ability of the NMR-based approach to adequately report on trends in bilayer elasticity.

### Frequency dependence of relaxation rates further informs the collective bilayer dynamics

The square-law relationship between the CH-bond order parameters and relaxation rates is measured at a particular Larmor frequency *v*_0_ (see Methods). To check whether the trend is dependent on *v*_0_ we used the simulation trajectories to analyze the CH-bond fluctuations for a range of Larmor frequencies. Figure 7 shows the results calculated at different *v*_0_ values for the DMPC bilayer in the liquid-disordered state. Regardless of the specific value of *v*_0_ the relaxation rates and squared order parameters are linearly dependent, illustrating that the square-law relationship holds for all the accessible frequencies, as seen experimentally (82). At lower frequencies the data are more spread out, while at higher frequencies the deviation from the best fit is smaller. There is also a systematic decrease in the slope with increasing frequency both as expected theoretically and seen experimentally (82). These results demonstrate that the trend encoded by these structural and dynamic aspects of the CH-bond fluctuations is not accidental, but rather arises from inherent bilayer elastic properties as they emerge from the collective atomic-level interactions of the lipids (see Discussion).

**FIGURE 7:**
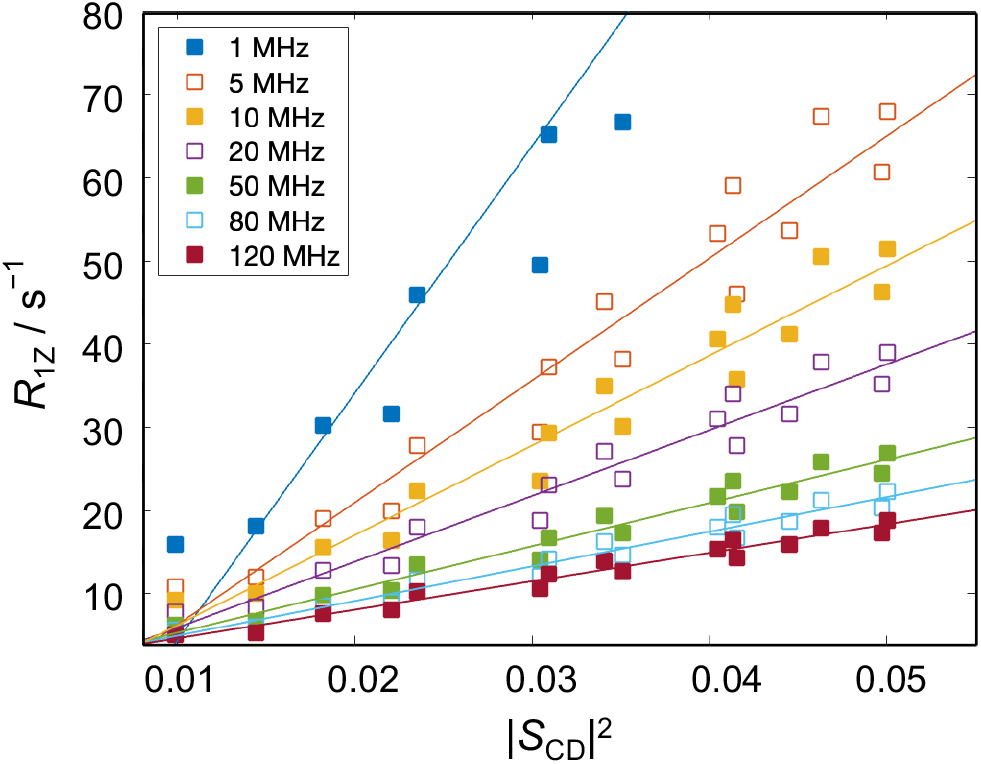
Frequency dependence of relaxation rates and order parameters for simulated bilayers. Shown are plots of the simulated ^2^H NMR relaxation rates of the acyl carbons of DMPC versus the squared order parameters at different Larmor frequencies *v*. Data are included for carbons on the *sn*-2 chain of DMPC in the cholesterol-free bilayer in the fluid state. Simulated results are depicted as squares and lines indicate the corresponding best fits. Note that the slope of the square-law increases with decreasing frequency consistent with experimental NMR data. The simulation was performed at 44 °C. **[1 column figure]**

## DISCUSSION

That the lipid composition of a membrane affects its structure and function is a universally acknowledged fact (9,10,83-86). However, considerably less is known about the level of cooperativity of the fluctuations of individual lipid molecules that give rise to the emerging bilayer properties. Despite the availability of an extensive amount of biophysical and NMR spectroscopy data, interpretations have remained controversial. We were motivated by the unique atomistic level information regarding both structure and dynamics offered by the NMR approach, as well as the ongoing need to continuously validate molecular dynamics (MD) force fields with experimental data (7,29). To this end, we used MD simulations to further explore the connection between lipid and bilayer dynamics, and its relation to NMR observables and bilayer elastic properties (5,38). Taking the unique features of the simulations into account, involving the fixed orientation of the bilayer normal, we derived a formulation that allows the simulation results to be directly compared to NMR measurements (34) of multilamellar vesicles (MLVs) as well as small unilamellar vesicles (SUVs) where the bilayer normal can adopt any angle with respect to the laboratory frame/magnetic field. Analysis of the simulation trajectories established the nematic-like behavior of the localized dynamics of the membrane lipids as suggested by NMR, namely their segmental 3D nature modulated by collective lipid dynamics. We saw a clear square-law relationship between the average order parameters of the lipid acyl carbon atoms and their respective relaxation rates, in excellent agreement with NMR results (35,82). The atomic resolution of the simulations allowed us to further identify the region in the bilayer where the square-law dependence holds, revealing the importance of interleaflet interactions for the collective dynamics of the lipids. The apparent bilayer bending moduli emerging from the application of an NMR-based formalism to the simulation data are corroborated by results from an alternative computational method, as well as NMR and neutron spin echo (NSE) experiments (85,86). Our MD simulations thus successfully replicate NMR observables and validate their interpretation while offering new insights about their physical origins.

### Theoretical framework for comparing molecular dynamics simulations to NMR measurements

There have been numerous occasions where results from MD simulations of lipid bilayers have been compared to NMR observables, such as acyl chain order parameters of the lipids or relaxation rates due to the underlying fluctuations (7,29,37,39,87). In the case of NMR, the elementary processes occur with correlation times near 1*/v*_0_ ≈ 10 ns, even though the relaxation times are in the 10–100 ms regime (weak collision limit) (34,56). Likewise atomistic MD simulations access the same simulation timescale even though the actual time needed to perform the simulation often differs by many orders of magnitude (so-called wall clock time) (9,59,87). Thus, both NMR spectroscopy and MD simulations entail measurements that are much longer than the actual processes of interest, in this case the lipid fluctuations, but the results of these measurements describe the same underlying dynamics. The order parameters depend entirely on a well-defined angle between the CH bond and the bilayer normal, and thus have a straightforward correspondence between the simulations and experiments. Still, the spin-lattice relaxation rate is not so trivial to translate between the two techniques. The problem arises because: (*i*) there are multiple frame transformations that need to be performed to relate the orientation of a lipid CH bond to the measured relaxation rate (see Fig. 4 in (57)), and (*ii*) NMR measurements are often performed on multilamellar lipid vesicles (MLVs) which have spherical symmetry, and as a result the bilayer normal can adopt any angle with respect to the magnetic field axis. These aspects reside in the liquid-crystalline nature of the membrane lipids and open up the question of what is the best way to encapsulate both the lipid structure and dynamics in a fully consistent manner?

The assumption of unrestricted isotropic motion circumvents the above problems and the resulting expression for the relaxation rate simplifies to a function dependent only on a single correlation time *τ*_C_ as shown in Eq. 9 and re-derived in the SM. In fact, the formula in Eq. 9 (and its equivalent for the carbon-13 relaxation rates) have been the ones used in comparisons of lipid dynamics between simulations and NMR in the past, see e.g., Refs. (32,71,72,88). In this solution NMR approach a spherical harmonics representation is often employed, where the CH bond fluctuations are described in terms of changes in their direction over time. The correlation function is thus written as:

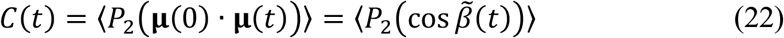

where *P*_2_ is the second-order Legendre polynomial (analogous to *S*_CD_ in Eq. 1). The CH bond orientation is defined by the scalar (dot) product of the unit vector **µ**(0) at time zero and the unit vector **µ**(*t*) at time *t* (Fig. 8 *A*). Fourier transformation of Eq. 22 gives *j*(*ω*) and *R*_1Z_ is subsequently calculated from the solution NMR result (Eqs. 9, S22, S23). Note that in the spherical harmonics representation, the pre-factor in Eq. 9 may look slightly different (see SM and Eqs. S22, S23). One should furthermore recall that Eq. 22 corresponds to the spherical harmonic addition theorem, i.e., where **μ**(0) · **μ**(*t*) = cos *β*(t) represents one side of a spherical triangle (Fig. 8 *A*) (89). The other sides are the CH bond orientation *θ*(0) (at *t*=0) and the bond orientation *θ*(*t*) (at time *t*), with |*ϕ*(*t*) − *ϕ*(0)| as the dihedral angle opposite to the 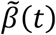 angle. Importantly, the angle 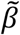 in Eq. 22 is different from the Euler angle *β* in Eqs. 2–4 because it does not depend on the director axis, or bilayer normal (Fig, 8 *A*). There is no projection axis (index) for the angular momentum, and hence Eq. 22 is applicable to solution NMR spectroscopy. For the case of solution NMR, the isotropic averaging is readily introduced leading to the results in textbooks. For multi-axis or composite motions, the orientational order parameters can be re-introduced leading to the so-called model-free approach for proteins in solution (90) as the limit of a generalized approach (35).

**FIGURE 8:**
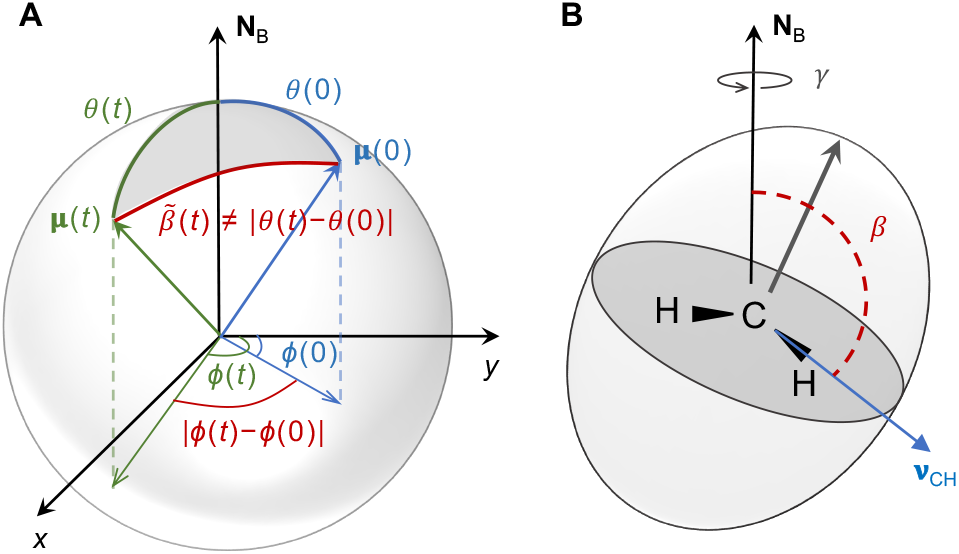
Schematic illustration of angles used in different approaches for analyzing CH bond fluctuations from MD simulations. (A) Spherical harmonic representation of the orientation of the CH bond at time = 0 and time = *t* described by unit vector **µ**(0) and unit vector **µ**(*t*). The corresponding spherical polar coordinates are the polar angles [*θ*(0), *ϕ*(0)]and [*θ*(*t*), *ϕ*(*t*)], respectively. The CH bond angle with respect to the bilayer normal **N**_B_ is *θ*(0) or *θ*(*t*). However, the angle *β*(*t*) whose fluctuations are analyzed with the use of the spherical harmonic addition theorem (Eq. 22) is different from |*θ*(*t*) − *θ*(0)| and does not depend on the director axis. (B) Alternatively, the orientation of the CH bond **v**_CH_ is described by the Euler angles (*α, β, γ*). Angles *β* and *γ* define the orientation of **v**_CH_ with respect to the bilayer normal **N**_B_ (director axis) and their fluctuations are analyzed in the new theoretical framework (Eq. 17). The schematic in (B) is adapted from Ref. (57). Note that use of the spherical harmonic addition theorem is applicable to isotropic liquids with unrestricted motions, while the representation in terms of Euler angles includes the director axis (bilayer normal) and orientational order parameters of the lipids. Analysis of multiscale composite motions of liquid-crystalline membranes thus becomes possible. **[1 column figure]**

On the other hand, lipid membranes are uniaxial liquid crystals in which the fluctuations are expressed in terms of orientational order parameters with respect to the bilayer normal, as shown in Fig. 8 *B* (see also Fig. 1 *A*) (35). Hence it is incorrect to apply the solution NMR limit. In an NMR experiment with liposomes, the bilayer normal can adopt all orientations with respect to the magnetic field and the CH bond fluctuations consequently might appear unrestricted. However, the CH bond fluctuations are not *isotropic* but rather *anisotropic* because of their inherent ordering with respect to the bilayer normal (i.e., their order parameters). That is why the analysis of solid-state NMR data assumes orientational averaging (the bilayer director can adopt all angles), but still retains information about the underlying ordering of the bonds (see SM). In simulations of bilayer patches, the bilayer normal is fixed but the CH bond dynamics are analogous to those in the liposomes measured experimentally with NMR spectroscopy. To connect the two approaches, we therefore derived an expression for the orientationally averaged relaxation rates, but in terms of the director frame, i.e., with respect to the local frame described by the bilayer normal (Eqs. 10 and 17). Stated differently, we take the simulation relaxation rates and then orientationally average them to compare directly to the experimental ones. Note that Eq. 10 for the orientationally averaged relaxation rate has an equivalent formulation in terms of spherical harmonics spectral densities (see Eq. S19). Importantly, the formulas presented here account for both the fixed bilayer normal in the simulations and the restricted anisotropic motion of the CH bond fluctuations described by segmental order parameters. They should thus be used instead of the solution NMR limit to compare simulation data to experimental NMR results.

To explicitly demonstrate the differences between the two theoretical approaches, i.e. using the spherical harmonic addition theorem with Eq. 22, versus the Wigner rotation matrix elements and Eqs. 16–17, we followed Ref. (72) to calculate the ^2^H NMR relaxation rates corresponding to the commonly used solution NMR limit. In particular, the spectral densities 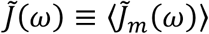 were obtained from the discrete one-sided Fourier transform of the correlation function using Eq. 22:

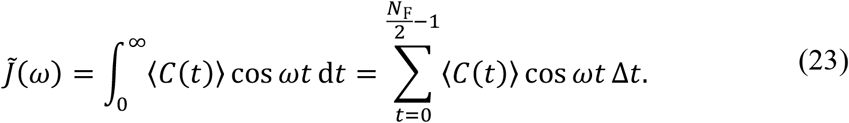

where ⟨*C*(*t*)⟩ is the orientational average of Eq. (22), i.e., the correlation function averaged over all bilayer orientations with respect to the laboratory frame. The relaxation rates were then calculated with Eq. 2.5 of Ref. (72) (also re-derived as Eq. S23 in SM):

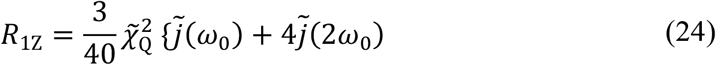

where the pre-factor in Eq. 24 is 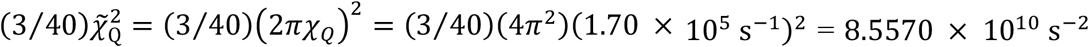. Accordingly, Fig. 9 shows the results from Eq. 24, noted as solution NMR, together with the results from Eq. 17, solid-state NMR and the experimental data (the latter two appear in Fig. 6 *A*). The solution NMR relaxation rates (filled red symbols) clearly deviate from the solid-state NMR data (filled green symbols), and the deviation is larger for the more ordered bilayer with 50% cholesterol. This result illustrates the underlying assumption behind Eq. 24 that the fluctuations are isotropic, so that there is no order in the system which is incorrect. Thus, when the order parameter is lower the two approaches are more alike, but when the order parameters are larger the differences become more pronounced.

**FIGURE 9:**
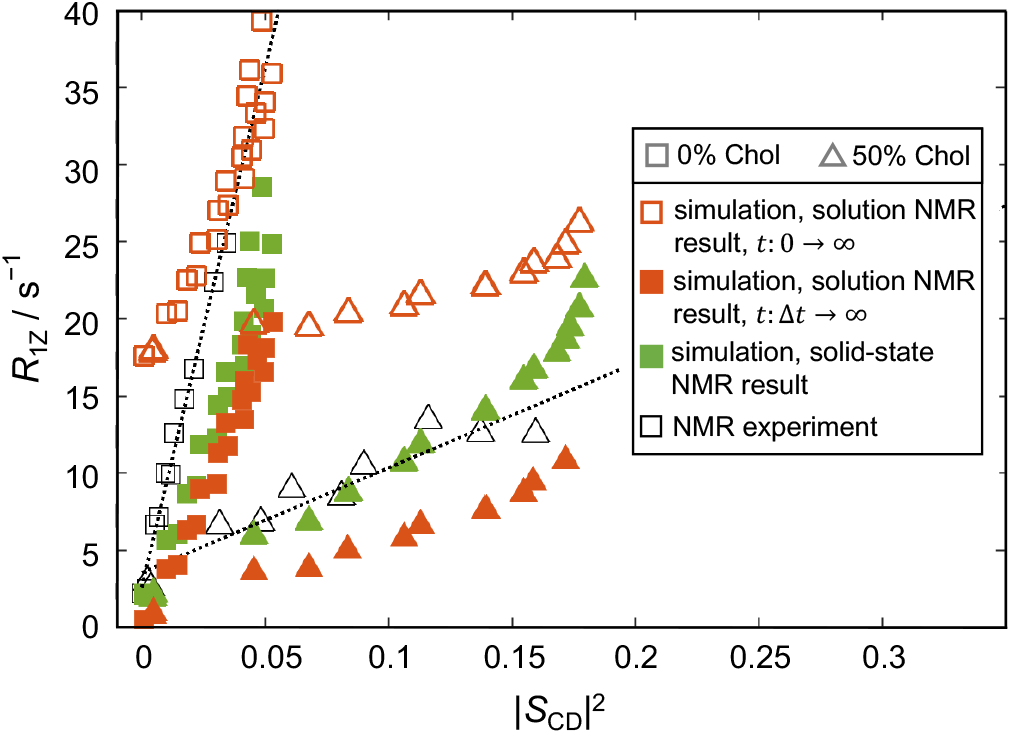
Comparison of relaxation rates from MD simulations calculated with different approaches. Shown are results from the solution NMR limit (Eq. 22–24) in red, the new framework (Eq. 17) in green, and experiments in black. Empty and filled red symbols correspond to the relaxation rates calculated from the discrete Fourier transform of the correlation function *C*(*t*) by integrating either from 0 → ∞ or from Δ*t* → ∞, respectively (Eq. 23). All simulations were performed at 44 °C. Quantitative agreement between the simulation and experimental relaxation rates requires consideration of both the sampling interval and whether solution NMR or solid-state NMR is used. See text for details. **[1 column figure]**

What is more, Fig. 9 shows also how the solution NMR approach suffers from the same sampling problem due to the discrete Fourier transformation as the solid-state NMR approach (see below). At *t* = 0 the correlation function from Eq. 22 is *C*(0) = 1 and when the integral in Eq. 23 goes from 0 to ∞, this results in a large shift in the spectral density and correspondingly, the relaxation rates (open red symbols). If instead the element at *t* = 0 is ignored, and the integral in Eq. 23 is evaluated from *t* = Δ*t* to ∞, the relaxation rates are closer in absolute value to the experiment. Note that the shift due to the *C*(0)Δ*t* term in Eq. 23 at *t* = 0 increases the magnitude of the relaxation rates but does not affect the slope of the square-law relationship, because *C*(0) = 1 for all carbons on the lipid chains. In contrast, in Eq. 15 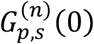 is the mean-square amplitude or variance of the Wigner rotation matrix elements, which is different for the various carbon atoms due to their inequivalent order parameters (Fig. S13). Consequently, ignoring the element at lag time = 0 in Eq. 15 would change not only the magnitude of the *R*_1*Z*_ values but also the slope of the square-law relationship. This aspect must be recognized in any validations of MD simulations or their force fields versus actual experimental NMR data.

### The nature of lipid dynamics in biomembranes

The spectral densities of the acyl chain CH bond fluctuations hold the key to the types of motions that the lipids exhibit while interacting with other lipids in a bilayer. It was recently shown that these spectral densities (calculated using the NMR solution limit described above) can be fit to a large sum of exponential functions (67). While the respective fits are very good, likely due to the large number of fitting parameters, the results point to an extensive hierarchy of motions with unclear origins, i.e., very fast dynamics modulated by slightly slower dynamics, modulated by yet slower dynamics, and so on. We show here that the spectral densities directly corresponding to the NMR measurements can instead be adequately fit to a simple power law function that has a clear physical interpretation pertaining to the nature of the underlying motions. The value of the single relevant parameter, the power exponent, indicates the degree of cooperativity of the lipid motions. For all carbons on all chains of all studied bilayers in the present study, we find the exponent to be around −1/2 (Fig. 2 *B* and Figs. S6–S8) consistent with collective segmental dynamics. This observation is further confirmed by comparison of the respective power exponents to those of local director (LD) vectors of varying lengths, whose orientations with respect to the bilayer normal fluctuate due to the thermal energy (Fig. 3, Figs. S10–S12). Regardless of which carbon the local director vectors originate from, their exponents decrease in an almost identical way with increasing vector length, illustrating the physical interpretation of the origins of this exponent: nematic-like motions at a value around – 1/2, and smectic-like dynamics at a value approaching −1 as the limit. Our results therefore are a direct manifestation of Occam’s razor and show how the seemingly complex interplay between lipid bond fluctuations is governed by a simple principle of segmental cooperativity.

### Quantum-mechanical view of the square-law dependence

Our simulations confirm the existence of the square-law dependence in simulated bilayers, in excellent agreement with NMR results for in vitro model lipid membranes. While we can distinguish two distinct regions of the lipid chains where this relationship holds and where it breaks (Figs. 4, 5, S14), the question remains as to the origins of this initially surprising functional correspondence between the order parameters and relaxation rates (35). Notably, the fluctuations of the CH bonds of a lipid reflect the changes in the energy of the atomic nuclei due to their magnetic or electric coupling to their surroundings. Such energy fluctuations result in transitions between different nuclear spin energy levels. The main energy levels of a nucleus in an NMR experiment are determined by the applied magnetic field (Zeeman effect), where transitions between these main energy levels (whose rate is quantified by *R*_1Z_) are enabled by additional energy perturbations due to the interactions of the nucleus with its surroundings. Weak coupling to the surroundings results in slower and less efficient relaxation while strong coupling gives rise to much faster relaxation rates. Thus, the rate of the relaxation depends on the strength of the energy perturbations near the resonance frequency. The transition probability between two main energy states is found to be proportional to the product of the squared matrix element of the perturbation and the density of states, according to Fermi’s golden rule. In the case of the fluctuating CH bonds, the matrix element is analogous to the mean-squared amplitude of the fluctuations, i.e., corresponding to the square of the segmental order parameter. Together with the square-law dependence, the NMR data show an additional dependence on inverse frequency in the form of a factor of *v* ^−1/2)^ that multiplies or scales the squared order parameter (that is why this term appears in Eq. 19). Because the latter is analogous to the density of states in the original formulation, the square-law relationship becomes a direct manifestation of time-dependent perturbation theory and Fermi’s golden rule in spectroscopy.

### Bilayer elastic properties emerging from the region of interleaflet contact

The existence of the square-law dependence shows clearly that the motional rates of the lipid bond fluctuations are *functionally* related to their mean-squared amplitudes, that is their relationship yields an elastic constant of the bilayer in a specific part of the lipid chains. These collective nematic-like dynamics at the region of direct contact between the two leaflets are closely related to the bilayer bending elasticity (Fig. 6). This observation is intriguing given the recently reported relationship between thickness fluctuations in that same bilayer region and the monolayer area compressibility modulus (70). In particular, extensive analysis of a large set of simulated lipid bilayers revealed a strong correlation in the height fluctuations of all carbon atoms outside of this region, suggesting a direct influence of interfacial tension (occurring at the water-hydrocarbon interface) on their dynamics (Fig. 1 in Ref. (70)). In contrast, the height fluctuations of the carbon atoms inside the region of leaflet-leaflet contact were decoupled from the interfacial dynamics, and hence the thickness fluctuations of that particular bilayer slab yielded the compressibility moduli of the two leaflets and the bilayer (70).

Emerging from the fluctuations in thickness and lipid chain CH-bond fluctuations in this middle bilayer region, the elastic moduli of both bending and compressibility suggest an intricate relationship between non-interfacial segmental dynamics modulated by interleaflet coupling and the energy required to bend or stretch the bilayer. These mechanical constants have been traditionally obtained from global bilayer properties governed by interfacial dynamics (e.g., thermal vesicle shape fluctuations or response to induced out-of-equilibrium deformations) in terms of underlying theoretical concepts of membrane elasticity (5,38,43,44,91). Instead, our results point to an alternative and underappreciated contribution from the more localized lipid dynamics at the bilayer midplane which appear to play a major role in the observed elastic behavior. Both approaches at the global and more local scales, have been shown to yield similar results for some bilayers (e.g., saturated lipids with and without cholesterol), but contradictory results for other types of bilayers (e.g., di-unsaturated lipids with cholesterol (86)). The sources of these discrepancies may lie in the time- and length-scales of the underlying dynamics (85,86) and are the subject of ongoing research.

### Validation of molecular dynamics simulations with experimental NMR data

As discussed above, proper comparison between the relaxation rates obtained from NMR and calculated from MD simulations requires special considerations which are taken into account in the theoretical framework presented here. The new formulation allows us to comment on a few key points pertaining to the relationship between theoretical MD simulations and experimental NMR relaxation studies. Here we recall that the simulation data are always discrete in nature, that is atomic coordinates are collected at some fixed time sampling interval (Δ*t* > 0). As a result, calculation of the spectral densities is inevitably done via a discrete (one-sided) Fourier transform of the correlation function (Eq. 15). This discrete transformation as illustrated in Eq. 15 produces a constant equal to the product of the autocorrelation at lag time = 0 (which is the variance of the data, var 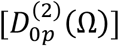 from Eq. 7) and the Δ*t* sampling interval. This constant is added to the spectral density at every frequency, and consequently it shifts the calculated relaxation rates in a Δ*t*-dependent manner which is an inevitable result of the discrete Fourier transform. Theoretically, the spectral density goes to infinite frequency, and its integral is equal to var 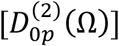 as can also be obtained from the second- and fourth-order Legendre polynomials (⟨*P*_2_⟩ and ⟨*P*_4_⟩) via a Clebsch-Gordan series expansion (see Table V in Ref. (74)). Equivalently, in the Fourier time domain, the correlation function goes to zero Δ*t* and the initial value equals var 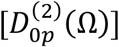 likewise. However, that limit is impossible to reach in simulations or experiments, and therefore we can only access a small window or bandwidth of the spectral density in the Fourier space.

The above notwithstanding we managed to alleviate the dependence on the discrete sampling interval Δ*t* by taking advantage of the fact that the correlation function of the CH bond fluctuations, expressed in terms of the second-rank Wigner rotation matrix elements, follows a power law function. As explained in Methods, this allowed us to fit the correlation function and use the best fit to resample it at smaller time intervals. For that, we found the smallest Δ*t*_fit_ that preserved the smoothness of the correlation function for each carbon, as shown in the Supplementary Material (Fig. S5). These Δ*t*_fit_ values correspond to the fastest rotations of the CH bonds that contribute to the spectral density, and are in the range of 0.5–30 ps. The results are in excellent agreement with the 5–20-ps correlation times predicted to arise from the bilayer *microviscosity* and account for the apparent enhancement in the relaxation of lipid bilayers relative to simple hydrocarbon fluids as measured with NMR spectroscopy (18). If a larger sampling timestep (such as the 40-ps output frequency in our simulations) were used, it would effectively mask these rapid bond rotations, while using an output frequency in the fs regime would be impractical for long MD simulations. Mathematical tricks for resampling of the correlation function are thus necessary to help bridge the gap between the discrete and continuum representations, even though an exact correspondence can never be achieved.

Another important observation from our analysis is that the square-law dependence can be used in a model-free way as illustrated by Fig. 6 *A*, to validate lipid force fields against NMR relaxation data. While the order parameter contains information about the average structure of the lipid chains—and therefore provides a static picture of their conformation—the spin-lattice relaxation rates describe the CH bond dynamics that produce these conformations, thus complementing the structural data. Figure 10 shows a comparison between the theoretically simulated and experimental order parameter profiles. Although the DMPC lipids in the presence of 0, 20, and 33% cholesterol in the simulations are more ordered than in the experiment, theory and experiment are in remarkable agreement in that we see smaller slopes of their square-law dependences corresponding to higher effective bending rigidities of the simulated bilayers (Fig. 6 *B*). This feature is expected from the known dependence of *κ*_C_ on lipid packing (and consequently order): the more disordered the lipids the lower the *κ*_C_ value (59,86). Interestingly, the simulated bilayer with 50 % cholesterol has slightly lower order parameters than the NMR measurements, and its square-law slope matches closely the experimental one for carbons C10 through C14 (Fig. 6 *A*). Therefore, while the overall trends in the deviations from experimental data are similar across the order parameters and relaxation rates, there may be differences indicating the ability of the force fields to better capture the elasticity of the bilayer or its average structure, thus highlighting a path forward towards their improvements. For that, the validation of lipid dynamics in the form of CH bond relaxation rates, as measured with NMR, becomes critical.

**FIGURE 10:**
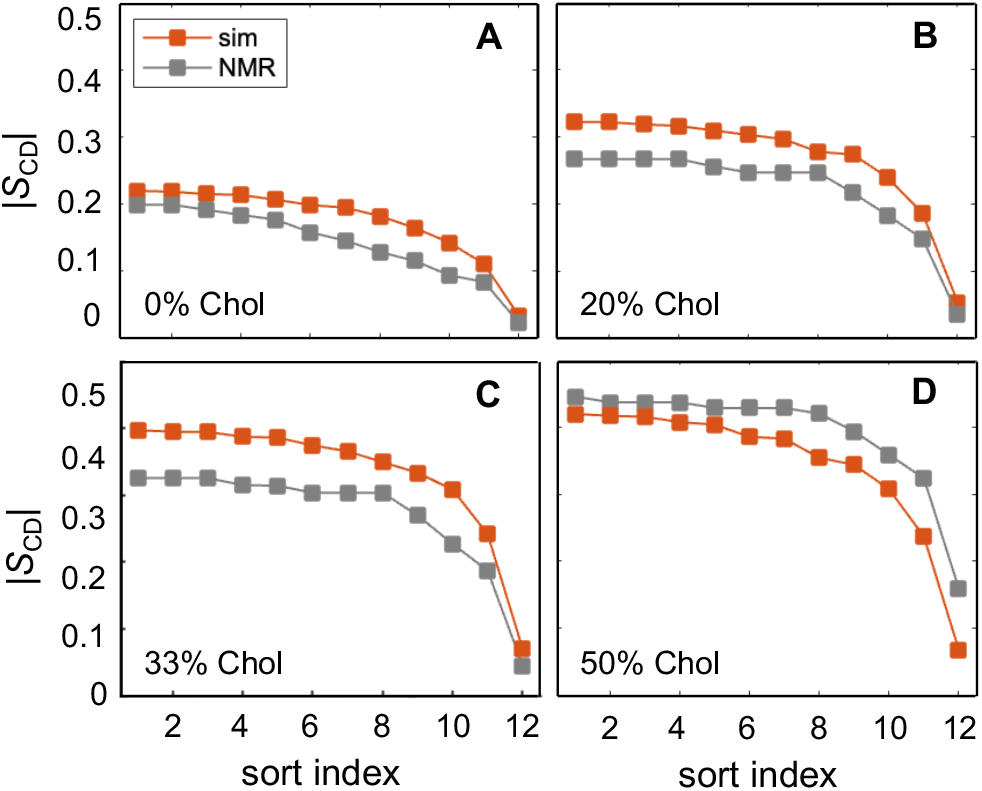
Comparison of DMPC order parameters obtained from simulations and NMR spectroscopy. Acyl chain order parameters are shown as a function of sort index (i.e., in descending order) for DMPC bilayers with 0, 20, 33, and 50% cholesterol either calculated from simulations (red) or reported from NMR (gray). Simulation data are average from the two chains. Experimental data were taken from Ref. (34) with the 0% cholesterol values interpolated from the data at 30, 50, and 60 °C. All simulations were performed at 44 °C. Validation of lipid force fields with NMR data requires consideration of both order parameters and relaxation rates. See text for details. **[1 column figure]**

Lastly, it is also worth noting that simulations are often validated against different types of experimental data, where differences in the types of samples and analysis employed by the various techniques may produce seemingly contradictory structural and/or dynamical features. One example is the recently reported problem in the lack of correspondence between the order parameters of sphingomyelin bilayers obtained from NMR and the bilayer structural properties (area per lipid and thickness) obtained from small-angle scattering techniques (6). Because the analysis of any experimental data adds another set of variables to its interpretation, a model-free comparison of the raw data is always optimal for a bias-free comparison. However, whenever there is a mismatch in the structure resolved by two approaches, the simulated system will inevitably fail to reproduce certain experimental observables.

## CONCLUSIONS

Experimentally observed relationships between the relaxation rates of CH bond fluctuations and their order parameters or the frequency at which they are measured have raised questions about the cooperativity of lipid motions in a bilayer and their link to membrane elasticity. We have shown herein that the local dynamics of the lipid acyl chains in the region of interleaflet contact as recapitulated by MD simulations are consistent with collective motions of the lipid segments analogous to those in nematic liquid crystals. For that, we developed a new theoretical framework that allows for direct comparison of the relaxation rates calculated from the simulations to those measured with NMR experiments, by accounting for the inherent differences between the two approaches. Following an NMR-based formalism for liquid-crystalline bilayers, we found that the square-law dependence between the rate and amplitude of the lipid CH bond fluctuations yields a frequency-independent estimate of an apparent bilayer bending modulus, which for DMPC bilayers follows the expected trends with addition of cholesterol. Our results are thus fully in line with NMR observables and their interpretation and offer an alternative protocol for extracting mechanical properties of the simulated membranes. Moving forward, it will be interesting to see whether our analysis is applicable to bilayers with mono and polyunsaturated lipids, as well as to more complex lipid mixtures aimed at establishing the universality of bilayer functional properties.

## AUTHOR CONTRIBUTIONS

M.D. and M.F.B. developed the concept and designed the research. G.K. produced the simulation trajectories and M.D. performed all computational analysis. R.A. performed the NSE experiments, including data collection and analysis. M.D., G.K., R.A., and M.F.B. wrote and edited the manuscript.

## DECLARATION OF INTERESTS

The authors declare no competing interests.

## ACKNOWLEDGMENTS

Klaus Gawrisch is acknowledged with gratitude for sharing his eternal insight and friendly collegiality over the years. We thank Steven Abel, Frederick Heberle, Kayla Sapp and Alexander Sodt for helpful advice and insightful discussions on the Fourier analysis of the time autocorrelation data from the simulations. We also gratefully acknowledge Andrew Erly and Trivikram Molugu for productive discussions on the interpretation of the simulation results. M.D. was supported by NIH F32GM134704. R.A. acknowledges NSF support through grant MCB-2137154. G.K. is supported by the HRH Prince Alwaleed Bin Talal Bin Abdulaziz Alsaud Institute of Computational Biomedicine at Weill Cornell Medical College through the 1923 Fund. M.F.B. acknowledges NIH support through grant R01EY026041 and NSF through grants MCB 1817862 and CHE 1904125. Access to the NGA-NSE beamline was provided by the Center for High Resolution Neutron Scattering, a partnership between the National Institute of Standards and Technology and the National Science Foundation under agreement no. DMR-1508249. Molecular dynamics simulations were performed using the Oak Ridge Leadership Computing Facility (Summit allocation BIP109) at the Oak Ridge National Laboratory (supported by the Office of Science of the U.S. Department of Energy under Contract No. DE-AC05-00OR22725).

## Supplemental Material

[Please see separate file]

## Notes

### Competing Interest Statement

The authors have declared no competing interest.

