## Supporting Material for "Molecular Simulations and NMR Reveal How Lipid Fluctuations Affect Membrane Mechanics"

The document includes 1 table, 15 figures and SI text.

The figures are as follows:


**Fig. S1** schematic of Euler and polar angles describing the forward and inverse transformations to and from the CH bond position

**Fig. S2–S4** correlation functions $G_{p,2}^{(n)}(t)$ for all carbons in the simulated bilayers with their corresponding best power-law fits

**Fig. S5** actual and resampled correlation functions $G_{0,2}^{(n)}(t)$

**Fig. S6** optimal $\Delta t_{fit}$ values (in ps) obtained from resampling of $G_{0,s}^{(n)}(t)$

**Fig. S7–S9** spectral density functions $J_{p,2}^{(n)}(\omega)$ for all carbons in the simulated bilayers with their corresponding best power-law fits

**Fig. S10** *b* exponents of the best power-law fits to the correlation functions $G_{p,s}^{(n)}(t)$ and spectral density functions $J_{p,s}^{(n)}(\omega)$ for all carbons in the simulated bilayers

**Fig. S11–S13** the spectral density functions $J_{0,2}^{(n)}(\omega)$ of local director vectors and their corresponding best power-law fits

**Fig. S14** bilayer structural properties (order parameters, lipid packing and thickness) as a function of Chol concentration

**Fig. S15** linear and log-log plots of $R_{1\mathrm{Z}}$ vs $|S_{\mathrm{CD}}|^2$ for all carbons in the simulated bilayers


**Table S1.** Full bilayer thicknesses calculated from the bilayer number density profile as described in Methods.

| Bilayer composition | Thickness / Å |
|---|---|
| DMPC | 51.2 |
| DMPC/Chol 80/20 | 55.2 |
| DMPC/Chol 67/33 | 56.4 |
| DMPC/Chol 50/50 | 55.6 |

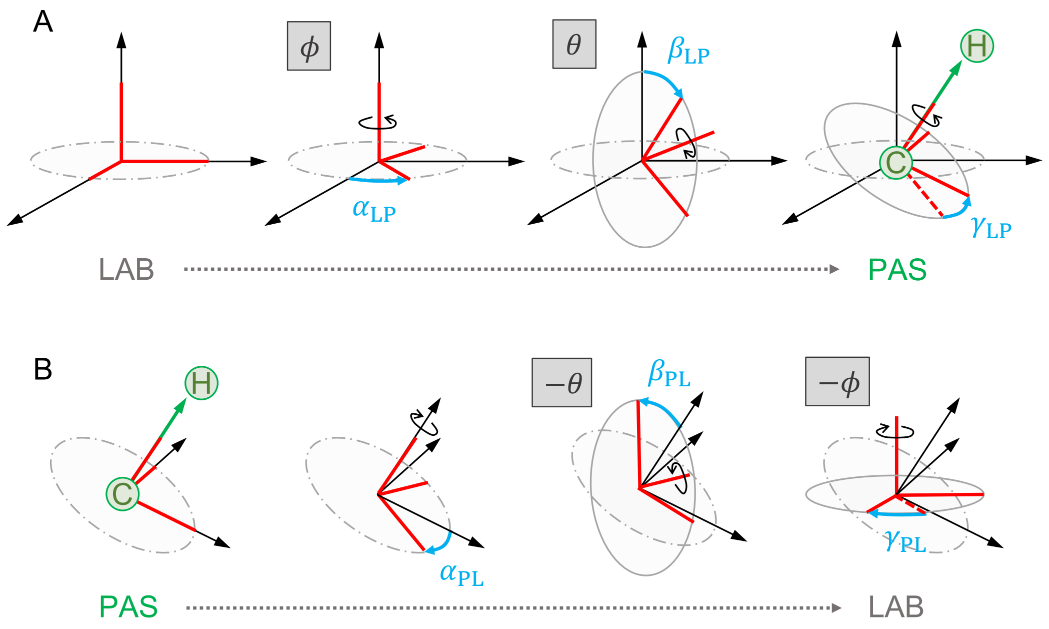


**Figure S1.** Schematic representation of the angles describing (A) the forward transformation from the laboratory frame (LAB) to the principal axis system (PAS) which is defined as the CH bond position (green), and (B) the inverse transformation from PAS to LAB. The Euler angles for the forward transformation are $\Omega_{LP} = (\alpha_{LP}, \beta_{LP}, \gamma_{LP})$ while the Euler angles for the inverse transformation are $\Omega_{PL} = (\alpha_{PL}, \beta_{PL}, \gamma_{PL}) = (-\gamma_{LP}, -\beta_{LP}, -\alpha_{LP})$. The corresponding polar angles $(\theta, \phi)$ are indicated in grey-shaded boxes next to their Euler angle equivalents.

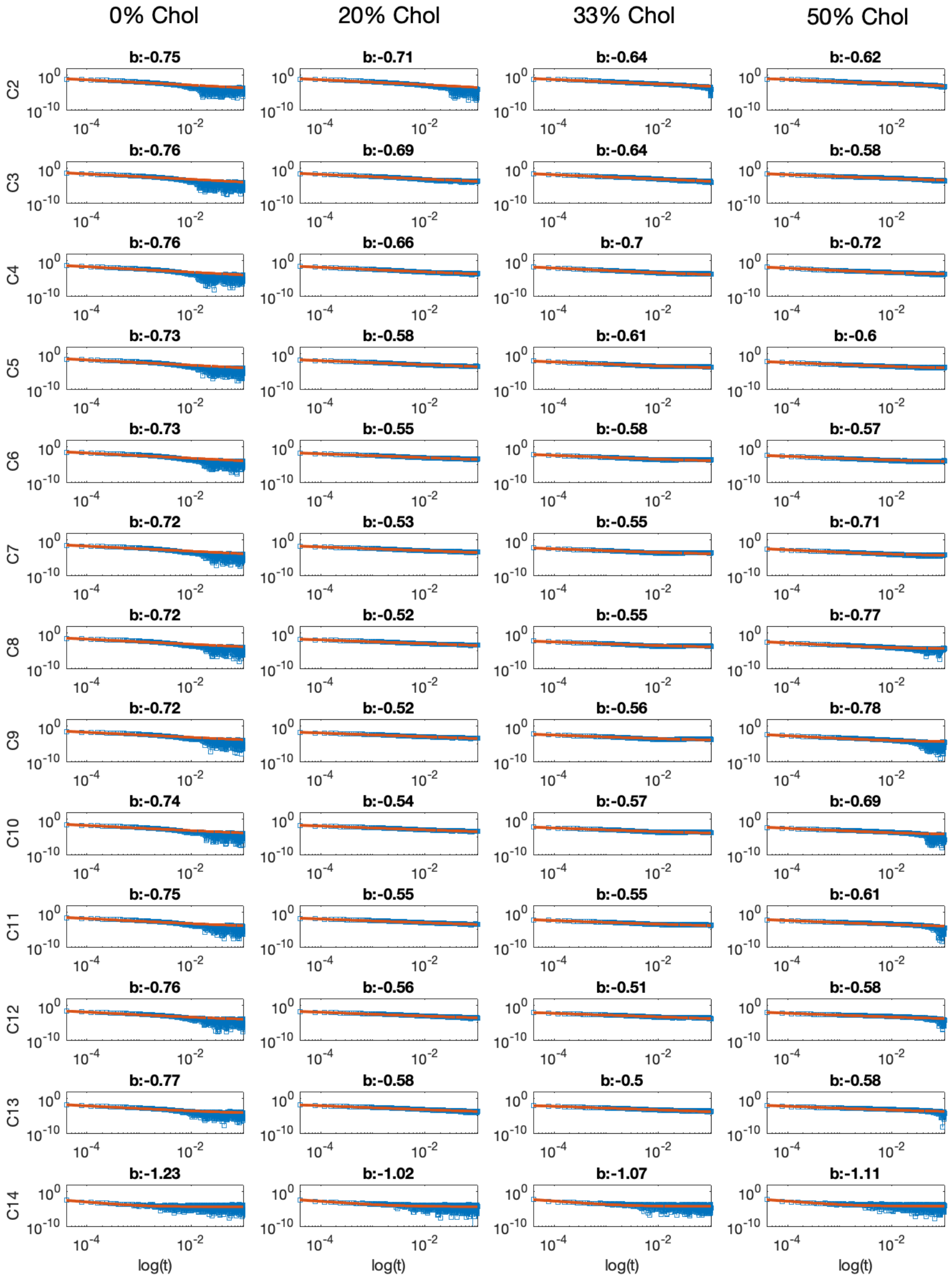


**Figure S2.** Log-log plot of the correlation function $G_{0,2}^{(n)}(t)$, $t \geq 1$ for each carbon atom on the sn-2 chain of DMPC in the bilayers with 0%, 20%, 33% and 50% Chol (blue data points), and corresponding best fit to a power-law function of the form $ax^b + c$ (red solid line). Shown above each plot is the $b$ exponent of the best fit. Only the first 100 ns of the correlation function are shown to better illustrate the fit. All plots have the same $x$- and $y$- axes.

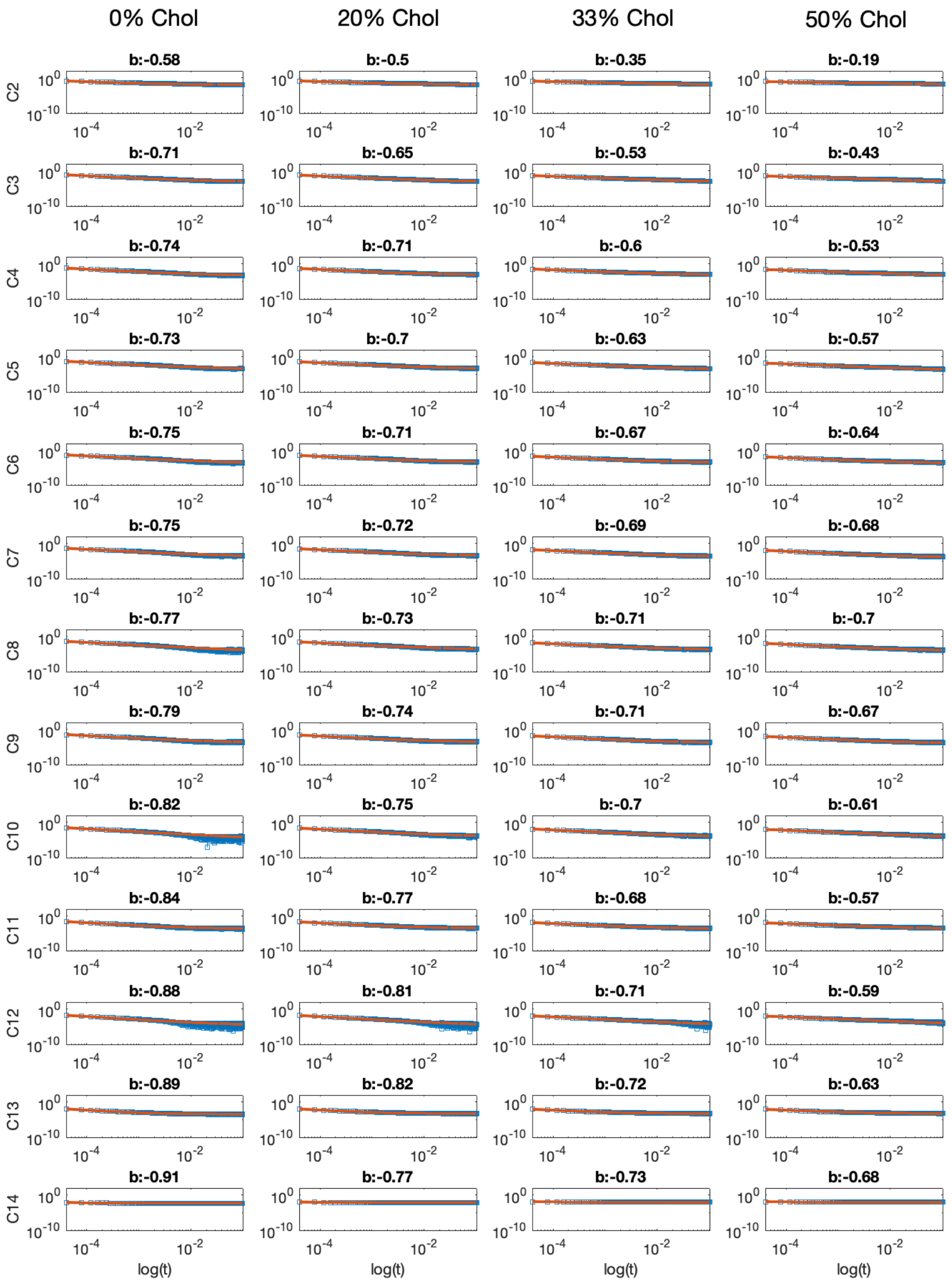


**Figure S3.** Log-log plot of the correlation function $G_{1,2}^{(n)}(t)$,$t \geq 1$ for each carbon atom on the sn-2 chain of DMPC in the bilayers with 0%, 20%, 33% and 50% Chol (blue data points), and corresponding best fit to a power-law function of the form $ax^b + c$ (red solid line). Shown above each plot is the $b$ exponent of the best fit. Only the first 100 ns of the correlation function are shown to better illustrate the fit. All plots have the same $x$- and $y$- axes.

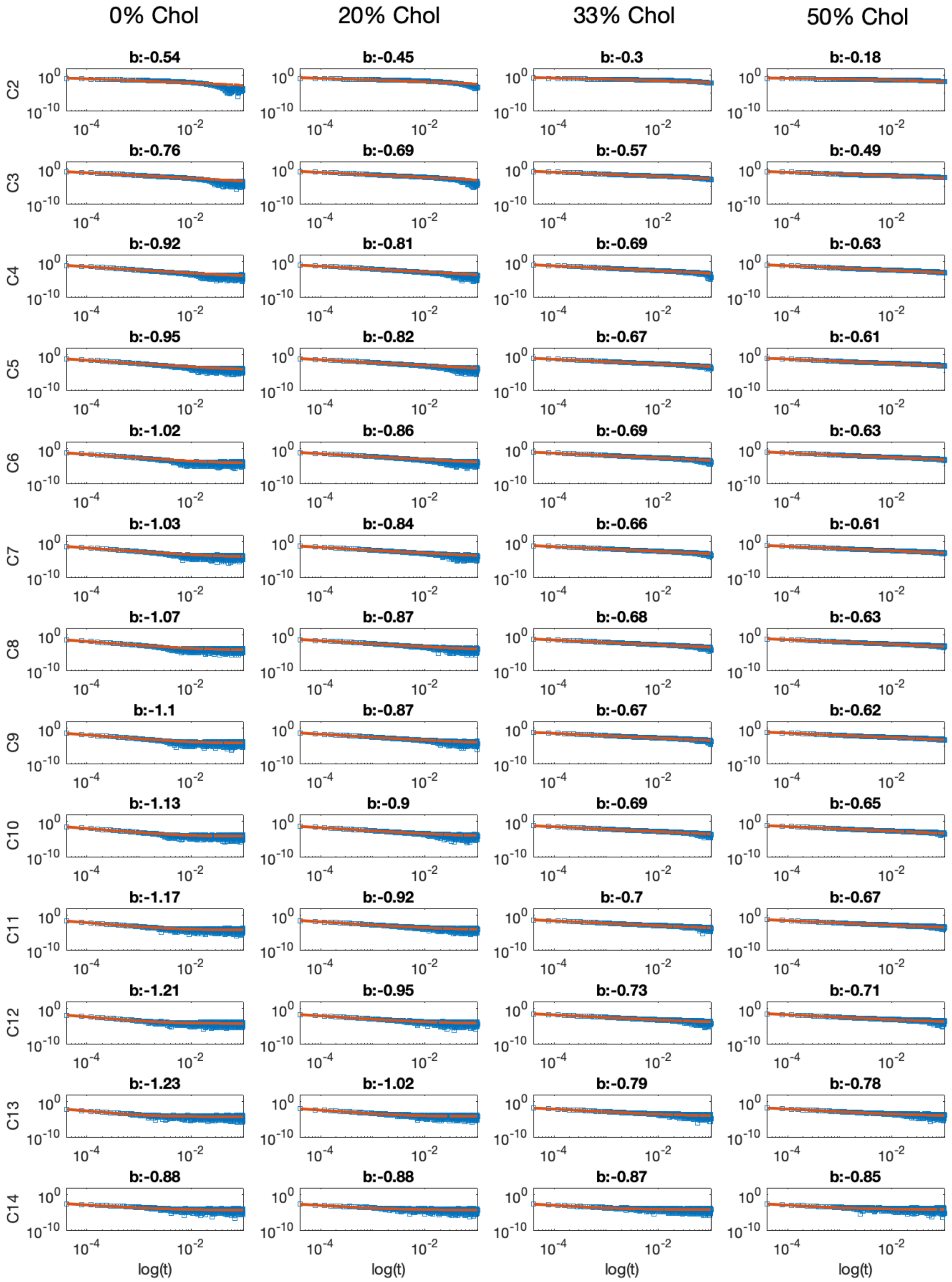


**Figure S4.** Log-log plot of the correlation function $G_{2,2}^{(n)}(t)$, $t \geq 1$ for each carbon atom on the sn-2 chain of DMPC in the bilayers with 0%, 20%, 33% and 50% Chol (blue data points), and corresponding best fit to a power-law function of the form $ax^b + c$ (red solid line). Shown above each plot is the $b$ exponent of the best fit. Only the first 100 ns of the correlation function are shown to better illustrate the fit. All plots have the same $x$- and $y$- axes.

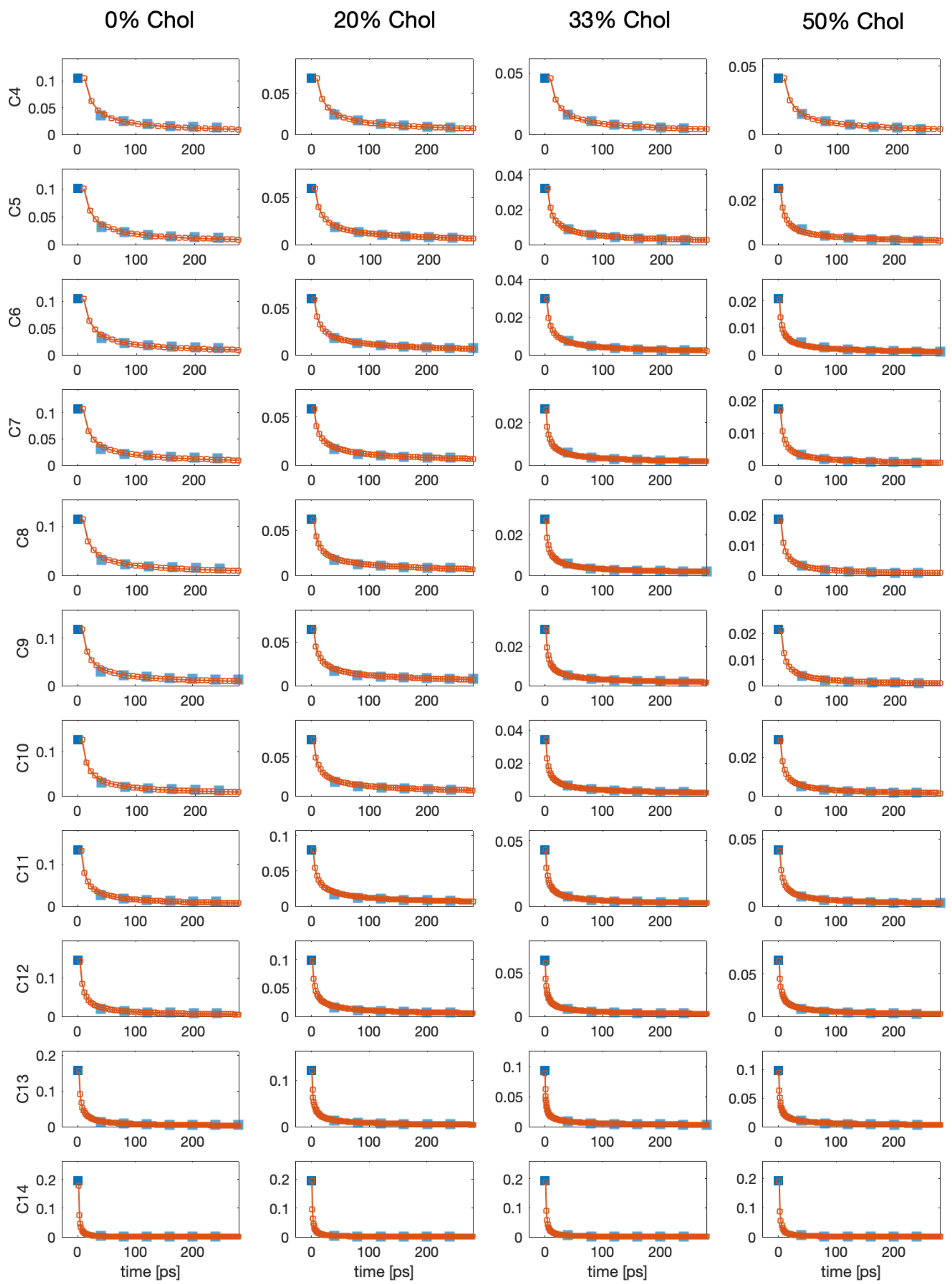


**Figure S5.** First 200 ps of the correlation function $G_{0,2}^{(n)}(t)$, $t \geq 0$ for carbons 4-14 on the sn-2 chain of DMPC in the bilayers with 0%, 20%, 33% and 50% Chol sampled at the original $\Delta t = 40$ ps (blue shaded squares), and the resampled correlation function from the best fits from Figure S2 sampled at the optimal $\Delta t_{fit} < \Delta t$ (red). $G_{0,2}^{(n)}(0)$ is highlighted in solid blue color.

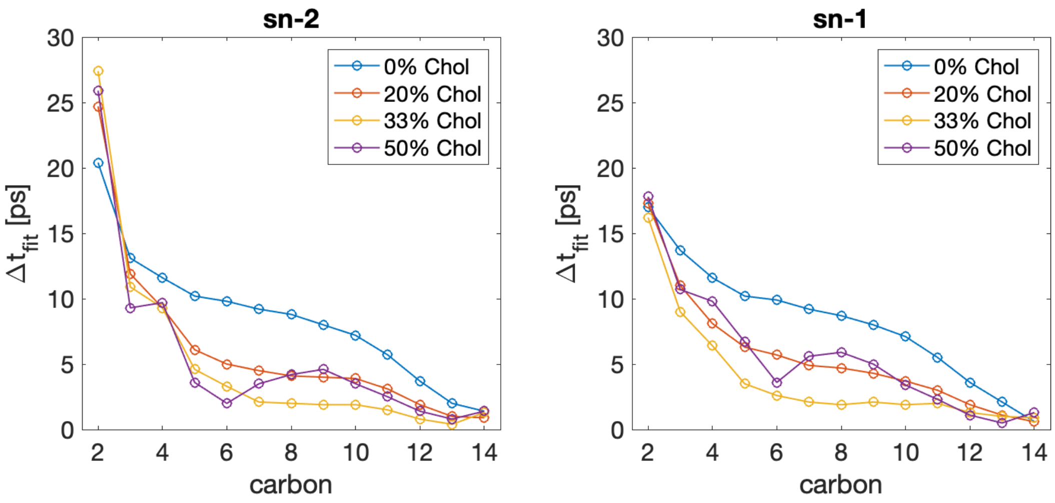


**Figure S6.** Optimal $\Delta t_{fit}$ values (in ps) obtained from resampling of $G_{0,s}^{(n)}(t)$ using the best fit (see Figure S5) for each carbon on the *sn*-1 and *sn*-2 chains of DMPC in the simulated bilayers. The simulation output frequency (and thus, original $\Delta t$) in the trajectories is 40 ps. The decrease of $\Delta t_{fit}$ further down the chain towards the terminal methyl (C14) is consistent with faster rapid motions closer to the bilayer midplane.

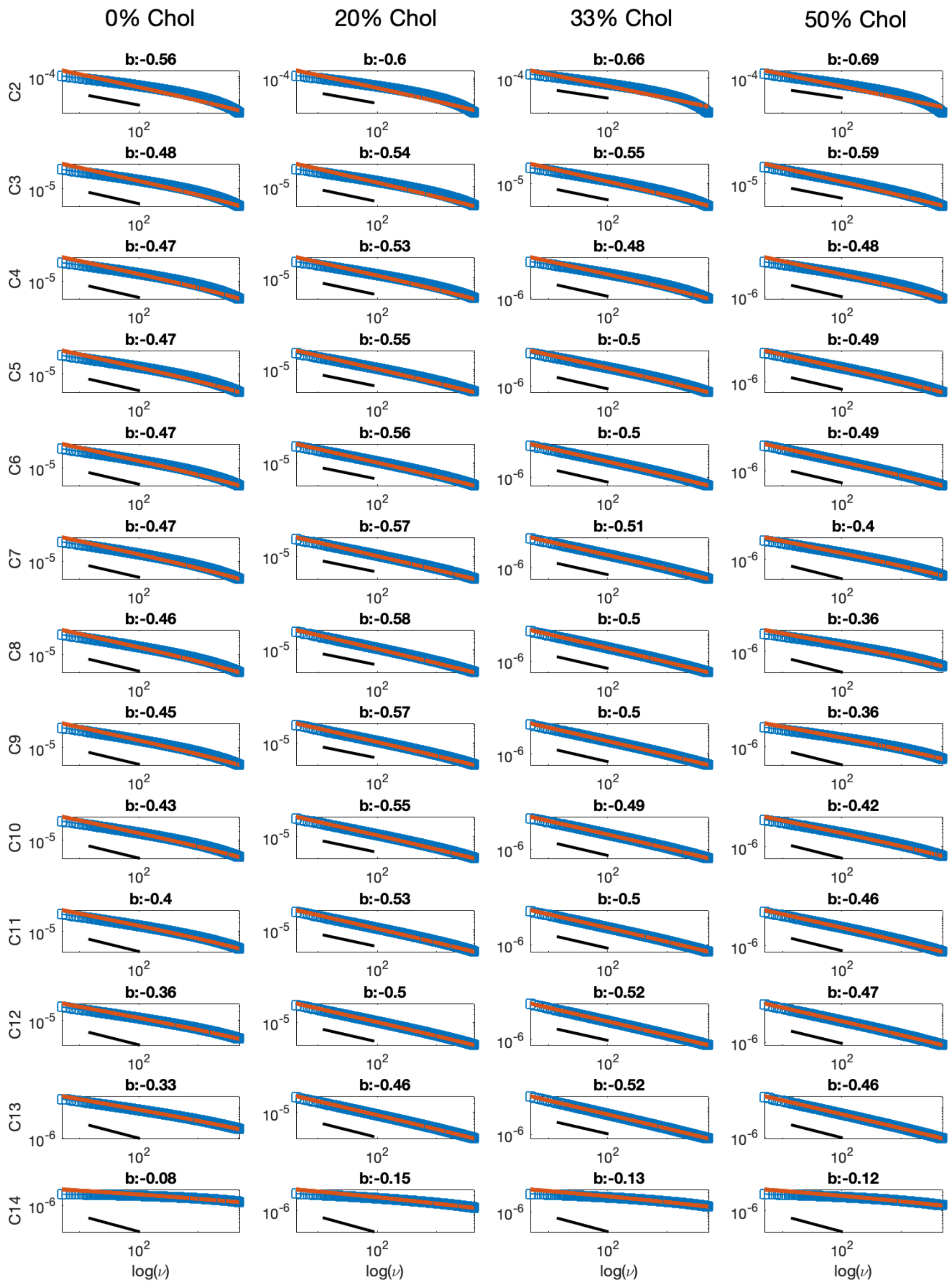


**Figure S7.** Log-log plots of the spectral density $J_{0,2}^{(n)}(\omega)$ (blue) for each carbon on the *sn*-2 chain of DMPC in the simulated bilayers, and its corresponding best fit to a power-law function of the form $ax^b + c$ (red solid line). Indicated above each plot is the $b$ exponent of the best fit. For visual comparison, a black line following a power-law with an exponent of −0.5 is shown below the data and fit.

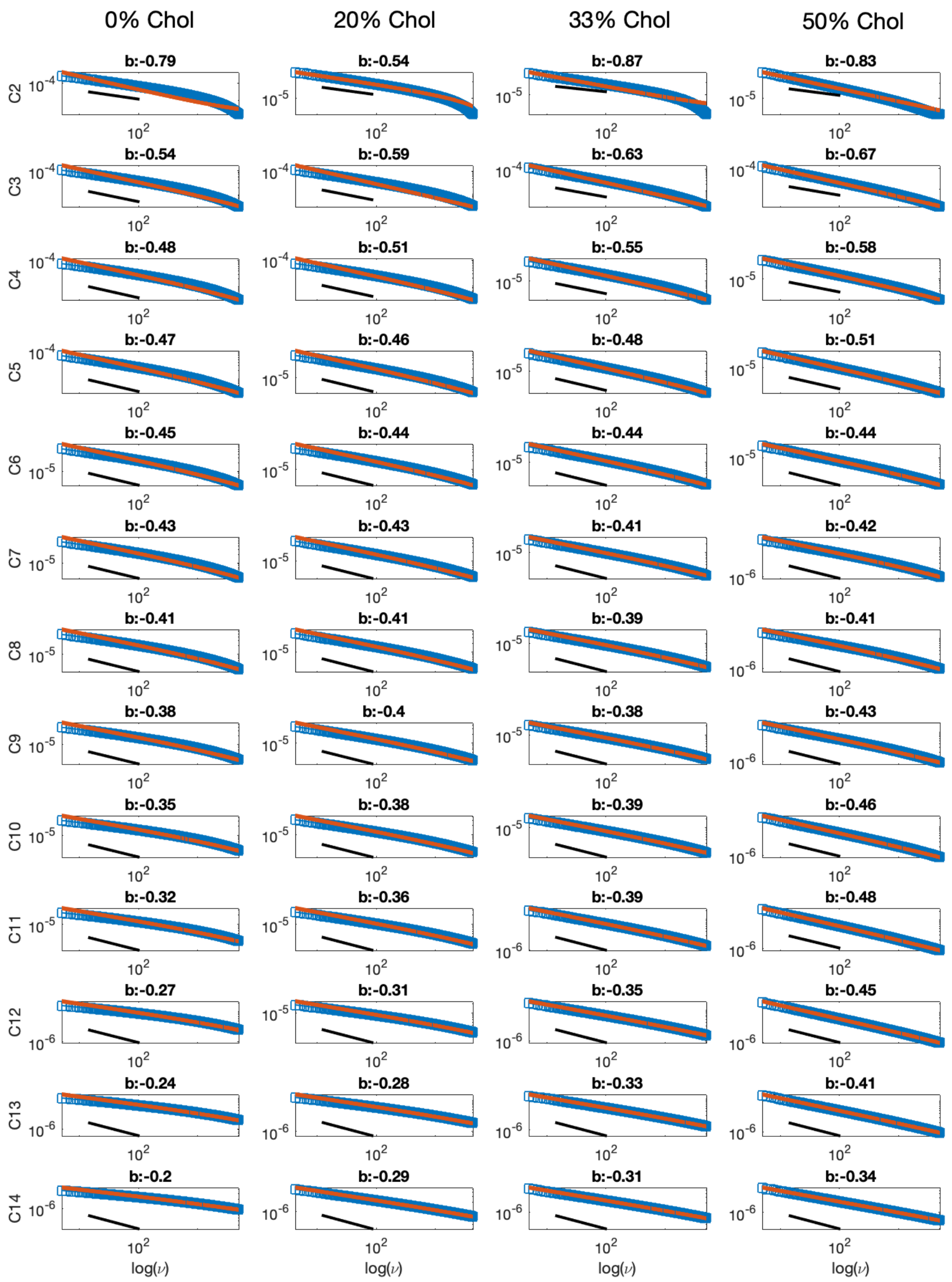


**Figure S8.** Log-log plots of the spectral density $J_{1,2}^{(n)}(\omega)$ (blue) for each carbon on the *sn*-2 chain of DMPC in the simulated bilayers, and its corresponding best fit to a power-law function of the form $ax^b + c$ (red solid line). Indicated above each plot is the $b$ exponent of the best fit. For visual comparison, a black line following a power-law with an exponent of −0.5 is shown below the data and fit.

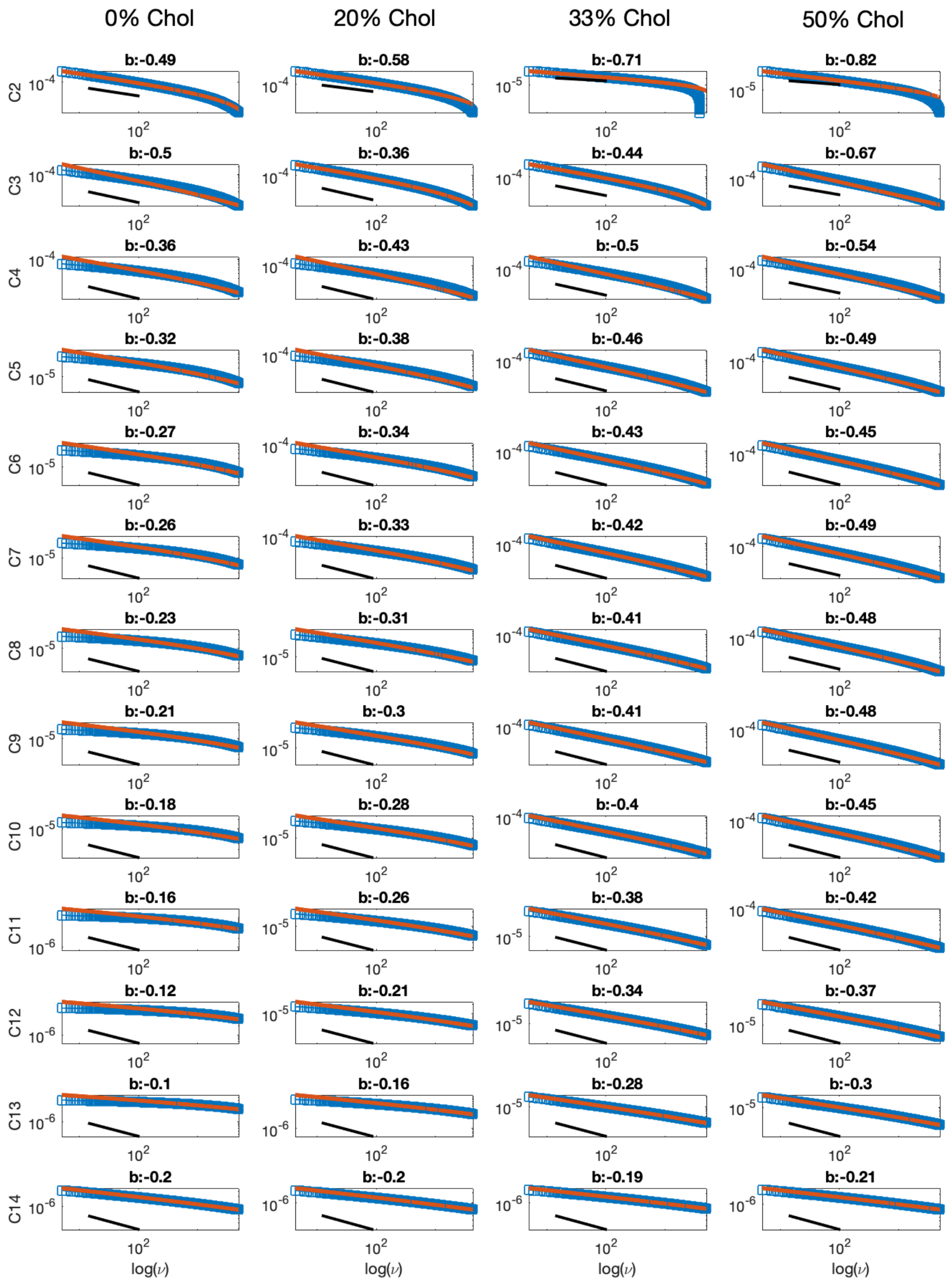


**Figure S9.** Log-log plots of the spectral density $J_{2,2}^{(n)}(\omega)$ (blue) for each carbon on the *sn*-2 chain of DMPC in the simulated bilayers, and its corresponding best fit to a power-law function of the form $ax^b + c$ (red solid line). Indicated above each plot is the $b$ exponent of the best fit. For visual comparison, a black line following a power-law with an exponent of −0.5 is shown below the data and fit.

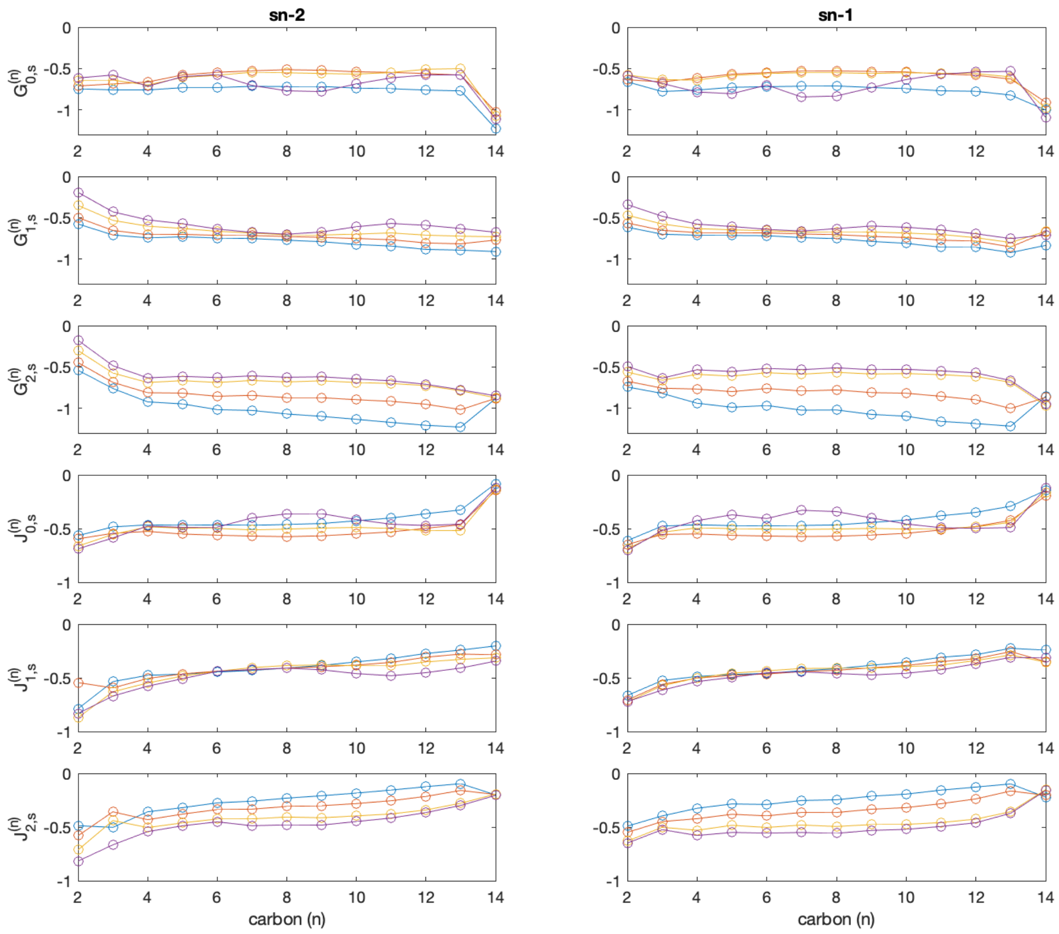


**Figure S10.** *b* exponents of the best fits to the correlation function $G^{(n)}_{p,s}(t)$ (top 3 rows) and spectral density $J^{(n)}_{p,s}(\omega)$ (bottom 3 rows) for each carbon on the *sn*-2 (left) and *sn*-1 (right) chains of DMPC in the bilayers with 0 (blue), 20 (red), 33 (yellow) and 50% (purple) cholesterol. Note that the $y$-axis for the correlation functions ranges from −1.3 to 0, while for the spectral densities it is between −1 and 0.

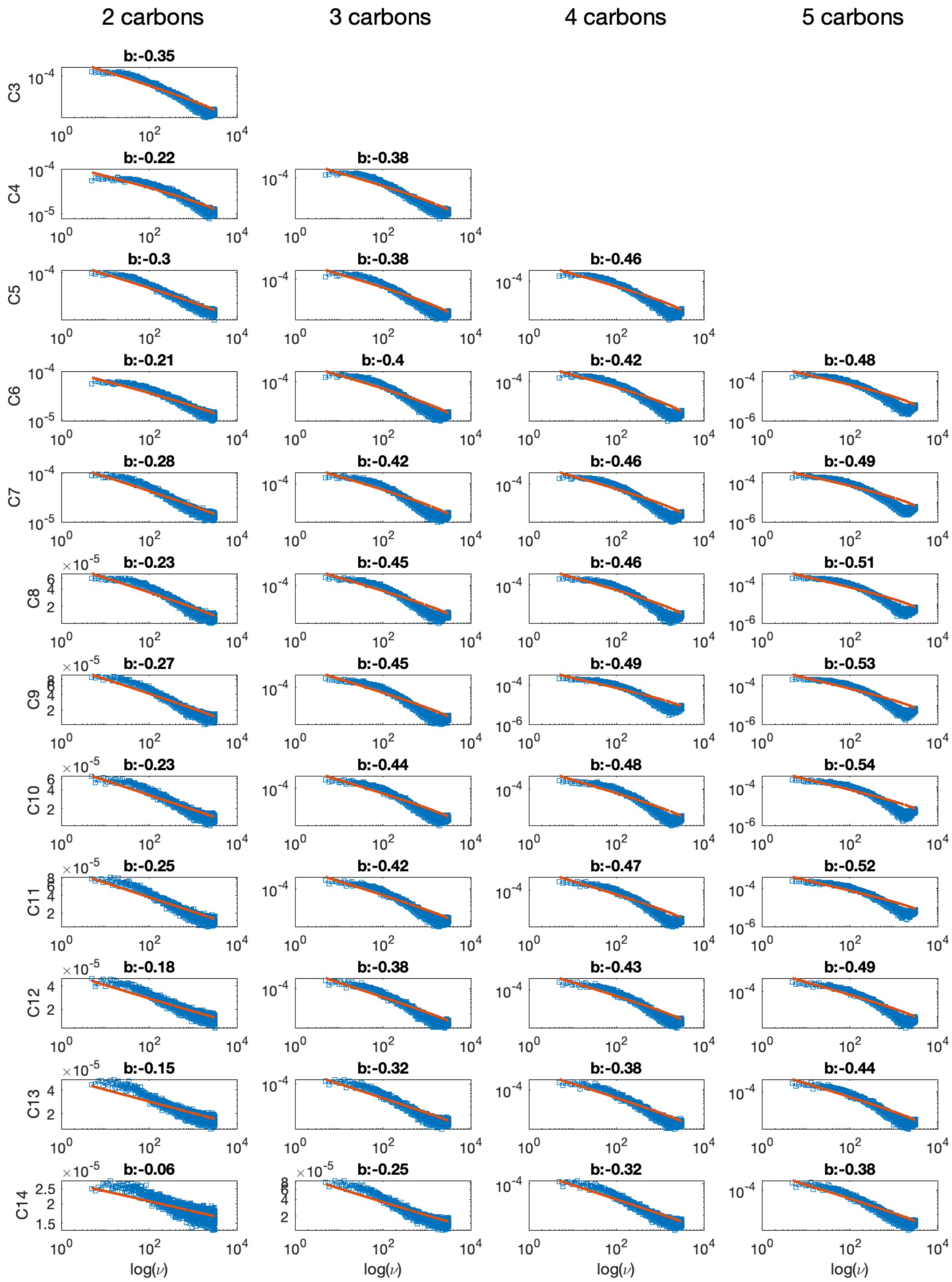


**Figure S11.** Log-log plots of the spectral density $J_{0,2}^{(n)}(\omega)$ of local director (LD) vectors of lengths 2–5 carbons. Each row corresponds to LD vectors originating from the same carbon (as indicated on the left). Raw data is shown in blue and best fit to a power law function of the form $ax^b + c$ is shown in red. The corresponding power coefficients of the best fits are indicated above the plots.

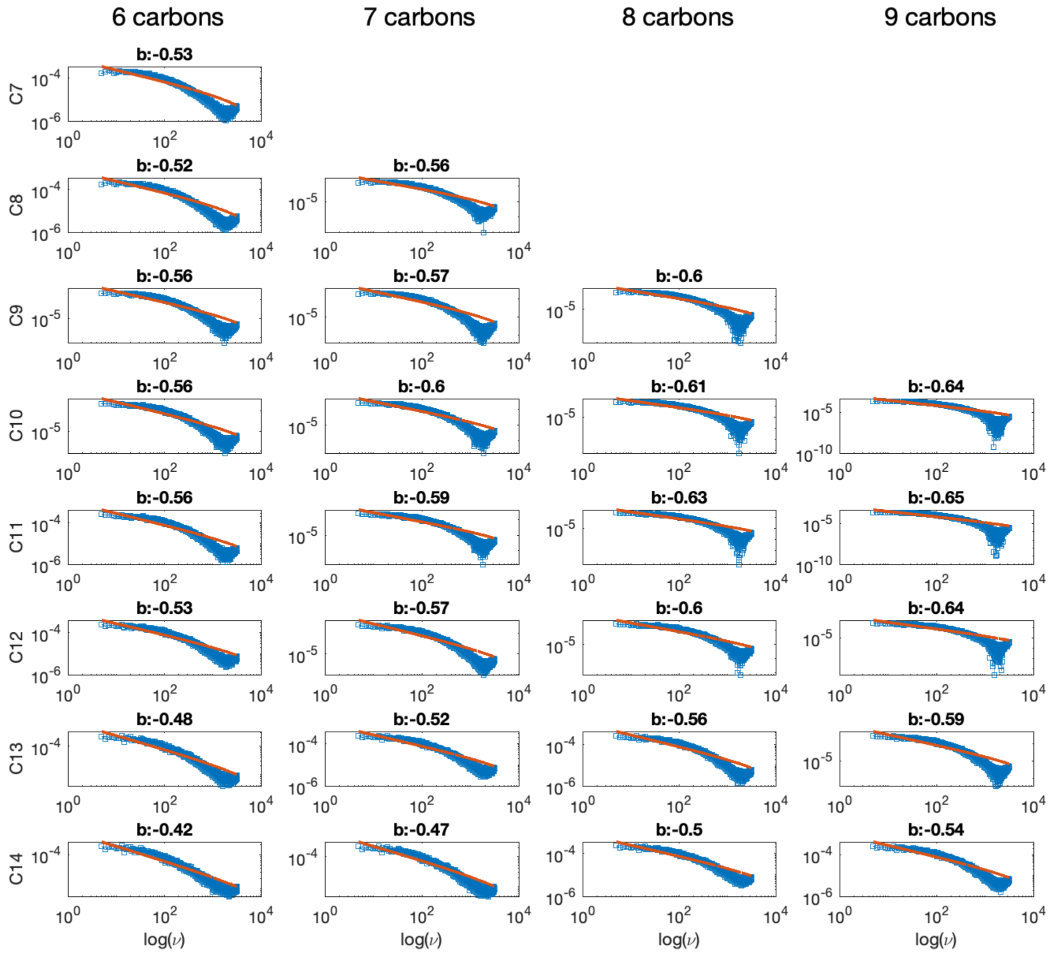


**Figure S12.** Log-log plots of the spectral density $J_{0,2}^{(n)}(\omega)$ of local director (LD) vectors of lengths 6–9 carbons. Each row corresponds to LD vectors originating from the same carbon (as indicated on the left). Raw data is shown in blue and best fit to a power law function of the form $ax^b + c$ is shown in red. The corresponding power coefficients of the best fits are indicated above the plots.

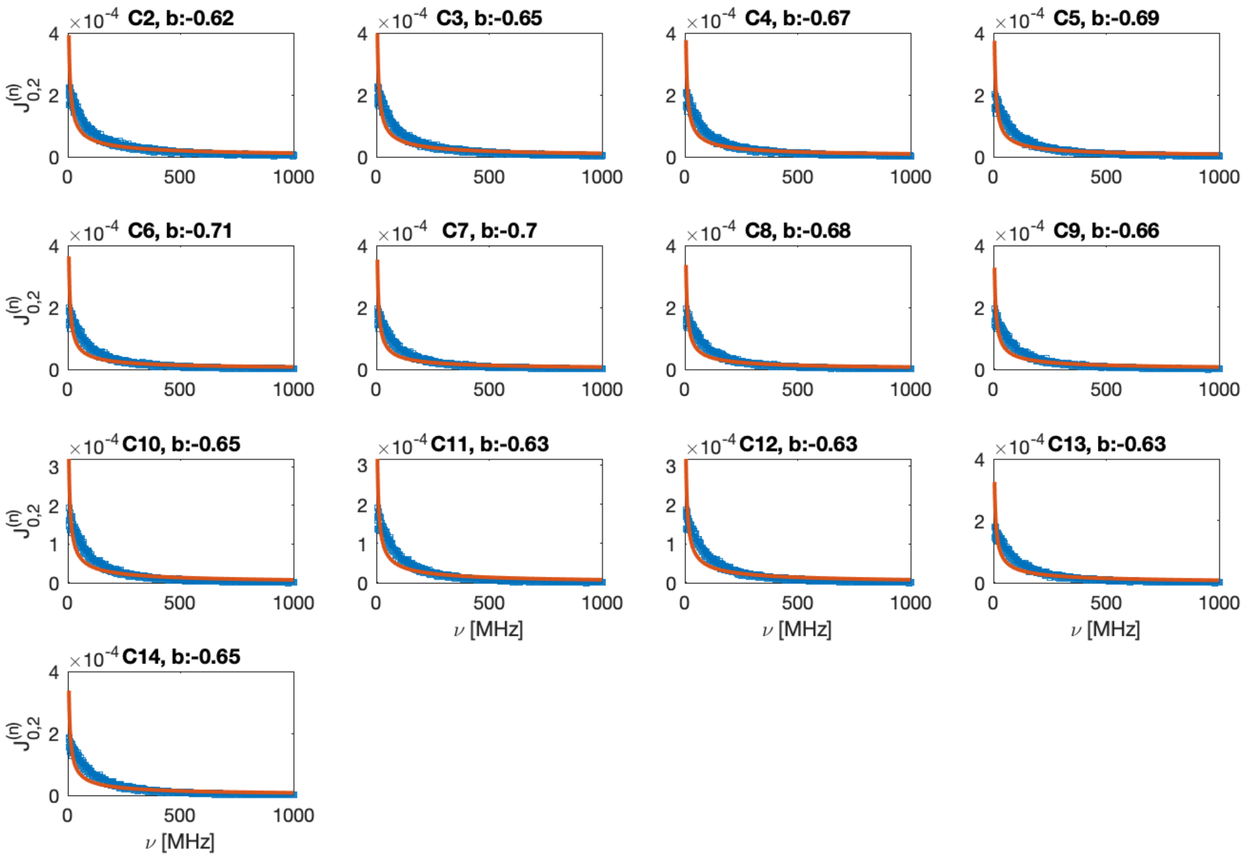


**Figure S13.** Log-log plots of the spectral density $J_{0,2}^{(n)}(\omega)$ of local director (LD) vectors connecting each carbon (as indicated above each plot) to the phosphate atom of the lipid. Raw data is shown in blue and best fit to a power law function of the form $ax^b + c$ is shown in red. The corresponding power coefficients of the best fits are indicated above the plots.

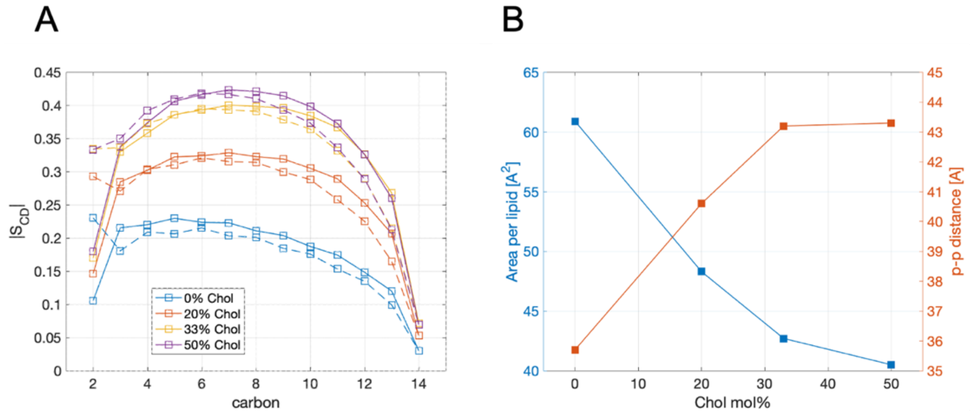


**Figure S14.** (A) Order parameter profiles for the two chains of DMPC, *sn*-1 (dashed) and *sn*-2 (solid), in the simulated bilayers. (B) Average area per lipid (blue, left axis) and phosphate-to-phosphate distance (red, right axis) as a function of cholesterol concentration.

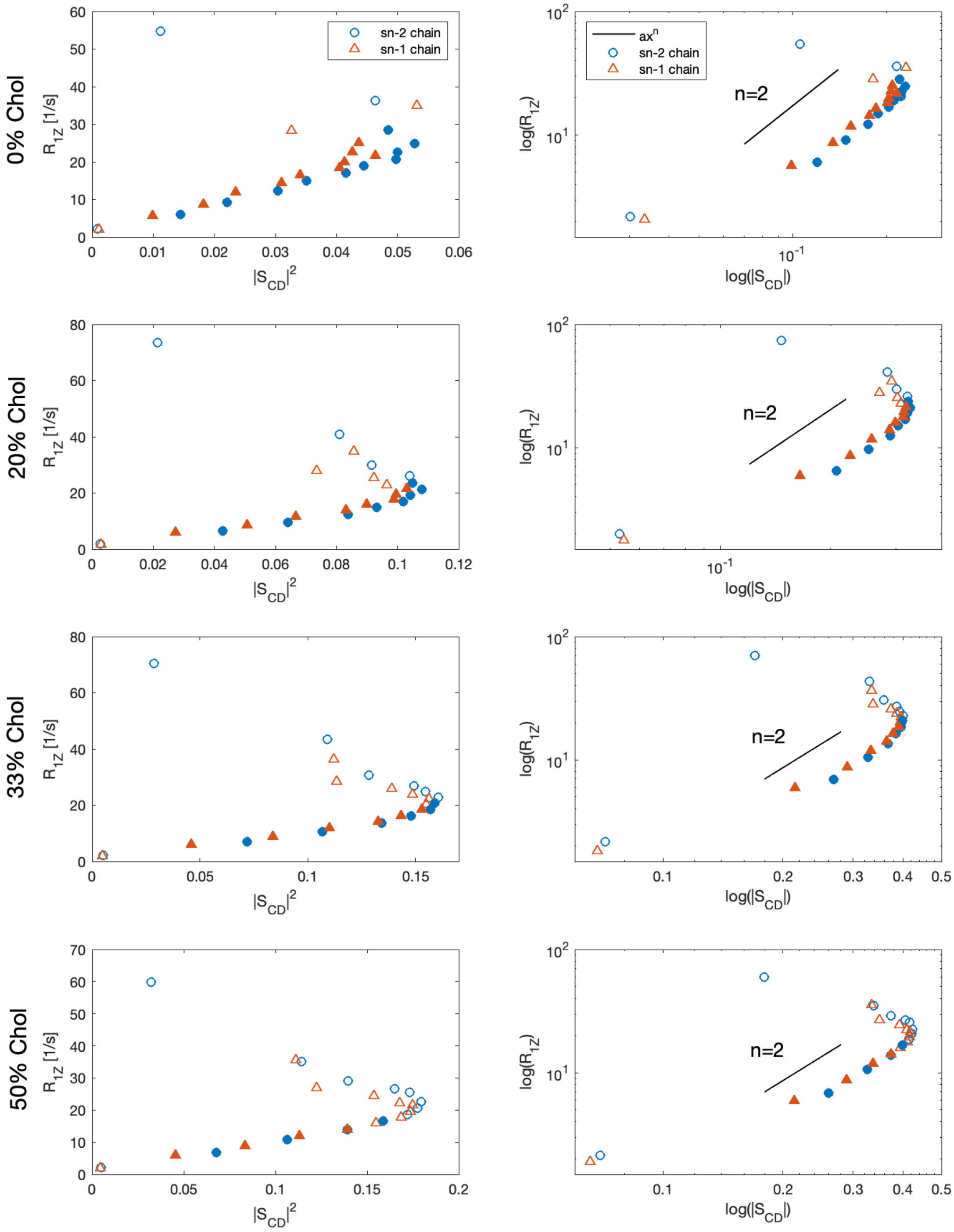


**Figure S15.** Relationship between spin-lattice relaxation rate $R_{1Z}$ and order parameter $|S_{\mathrm{CD}}|$ for all carbons of DMPC in the simulated bilayers. Carbons on the *sn*-2 chain are shown as blue circles and on the *sn*-1 chain as red triangles. Filled symbols indicate the carbons from each chain that follow the square-law relationship and were used in subsequent analysis of the apparent bilayer bending rigidity. On the left $R_{1Z}$ is shown as a function of $|S_{\mathrm{CD}}|^2$ while on the right a log-log plot illustrates the model-free relationship between $R_{1Z}$ and $|S_{\mathrm{CD}}|$. Also shown for comparison on these plots is a sample square-law function of the form $ax^n$ with $n = 2$ in black. Note that the $x$- and $y$- axes of the plots are purposefully different to better illustrate the spread in the data.

### Extended Methods

#### Neutron Spin Echo experiments

Suspensions of 100-nm unilamellar vesicles were prepared at a lipid concentration of 50 mg/mL, using standard vesicle extrusion methods (1). The samples were prepared by mixing protiated DMPC and cholesterol in chloroform at the required mole fractions. This step was followed by evaporation of chloroform under an inert gas stream and subsequent drying under vacuum overnight at 45 °C. The dry lipid films were then hydrated with a 10 mM deuterated sodium phosphate buffer (prepared with $D_2O$ instead of $H_2O$) at 45 °C with intermittent vortex mixing. The hydration of the lipid films at elevated temperature facilitates dispersion and promotes mixing of DMPC and cholesterol within the formed bilayers. The suspension was then subjected to at least 5 freeze/thaw cycles using a –80 °C freezer and a warm 45 °C water bath. The suspension was then extruded using an automated mini-extruder (Avanti Polar Lipids) by passing the suspension 31 times through a polycarbonate filter (pore size = 100 nm). The extruder setup was heated to 45 °C during all extrusions. After extrusion, the samples were kept in a Peltier box at 45 °C until measured.

The NSE experiments were conducted on the NG-A NSE spectrometer at the NIST Center for Neutron Research (NCNR) over a $q$-range of 0.04 $Å^{-1}$ to 0.1 $Å^{-1}$, where $q = 4\pi \sin\theta/\lambda$ is the wavevector transfer defined by the neutron wavelength, $\lambda$, and the scattering, $2\theta$, measured relative to the incident beam. Measurements of the instrumental resolution and deuterated buffer were performed under the same configuration for data reduction and normalization. Data reduction was performed using the Data Analysis and Visualization Environment (DAVE) software developed at NIST (2).

#### Theoretical background: nuclear spin relaxation of lipid membranes

In general, the NMR spectra and relaxation times are governed by the Hamiltonian for coupling of the nuclear spins to the environment of the molecule or material. The main Hamiltonian is due to the Zeeman interaction of the nuclear magnetic moment with the applied external (static) magnetic field and will not be further discussed here. The smaller perturbing Hamiltonian contains the information of interest to chemists or physicists. It is due to the chemical shift, the direct (through space) magnetic dipolar interaction, the indirect (through bond or electron-mediated, spin-spin) dipolar coupling, and the electric quadrupolar coupling in the case of nuclei with a quadrupolar moment describing the non-spherical nuclear charge distribution. The perturbation is typically considered within the quantum-mechanical interaction picture.

**Role of average Hamiltonian and correspondence to bilayer average structure.** When motion is present as in the case of liquid crystals, liquids, and even molecular solids, one must consider the average value of the perturbation and the fluctuating parts. The secular (time-independent part) part governs the lineshape while the non-secular (time-dependent part) part governs the relaxation. Notably, the fluctuations of the Hamiltonian occur with respect to the mean value so that the fluctuating part is given by: $\widehat{H}_\lambda{}' = \widehat{H}_\lambda - \langle \widehat{H}_\lambda \rangle$ , where $\lambda = \mathrm{Q}$ for the quadrupolar interaction in $^2$H NMR spectroscopy. The time-independent average Hamiltonian is secular; hence it commutes with the main Zeeman Hamiltonian and governs the NMR lineshape in accord with time-independent perturbation theory. Conversely, the averaged value of the Hamiltonian is non-zero and it must be subtracted to yield the fluctuating part that governs the relaxation. The fluctuating part is non-secular and affects the energy level transitions as treated by time-dependent perturbation theory.

**Correspondence to experimental relaxation rates in NMR spectroscopy.** Next, we apply time-dependent perturbation theory within the Redfield approximation (3). After transforming the principal values of the coupling (EFG) tensor from the molecule-fixed principal axis system (PAS) within the molecule to the laboratory frame, the irreducible correlation functions correspond directly to the Wigner rotation matrix elements. They read:

$$G_m^{\mathrm{lab}}(t) = \left\langle \left[D_{0m}^{(2)}(\Omega_{\mathrm{PL}};t) - \langle D_{0m}^{(2)}(\Omega_{\mathrm{PL}})\rangle\right]^* \left[D_{0m}^{(2)}(\Omega_{\mathrm{PL}};0) - \langle D_{0m}^{(2)}(\Omega_{\mathrm{PL}})\rangle\right]\right\rangle \qquad \text{(S1)}$$

where $D_{0m}^{(2)}(\Omega_{\mathrm{PL}})$ are the second-rank Wigner rotation matrix elements for the transformation for rotations from the molecule-fixed PAS to the laboratory frame in terms of Euler angles $\Omega_{\mathrm{PL}} \equiv (0, \beta_{\mathrm{PL}}, \gamma_{\mathrm{PL}})$ for transformation form the principal axis system (P) to the laboratory (L) frame (Fig. S1). The corresponding irreducible spectral densities of motion $J_m^{\mathrm{lab}}(\omega)$ are the Fourier transform partners of the $G_m^{\mathrm{lab}}(t)$ correlation functions:

$$J_m^{\mathrm{lab}}(\omega) \;=\; \mathrm{Re}\int_{-\infty}^{\infty} G_m^{\mathrm{lab}}(t)e^{-i\omega t}\,dt \qquad \text{(S2)}$$

In the above formula $J_m^{\mathrm{lab}}(\omega)$ is a two-sided Fourier transform of the correlation function with limits of $(-\infty, +\infty)$. For more details the reader is referred to Refs. (4,5).

The relaxation rates in NMR spectroscopy correspond to various linear combinations of the irreducible spectral densities of motions $J_m(\omega)$ of the coupling Hamiltonian. In the case of solid-state $^2$H NMR of lipid membranes, the spin-lattice relaxation rate ($R_{1\mathrm{Z}}$) and the quadrupolar order relaxation rate ($R_{1\mathrm{Q}}$) are of interest (3). The spin-lattice relaxation rate is given by:

$$R_{1\mathrm{Z}} = \tfrac{3}{4}\pi^2\chi_{\mathrm{Q}}^2\left[J_1^{\mathrm{lab}}(\omega_0) \;+\; 4J_2^{\mathrm{lab}}(2\omega_0)\right] \qquad \text{(S3)}$$

where $\chi_{\mathrm{Q}} \equiv e^2qQ/h$ = 170 kHz is the static quadrupolar coupling constant (6). The value of the numerical pre-factor is thus $(3/4)\pi^2(1.70\,x\,10^5)^2$ = $2.1392 \times 10^{11}\,s^{-2}$. The irreducible spectral densities of motion $J_m^{\mathrm{lab}}(\omega)$ are the Fourier transform partners of the $G_m^{\mathrm{lab}}(t)$ correlation functions (Eq. S1) and are directly connected to the observable relaxation rates in NMR spectroscopy.

**Formulation of spherical harmonic correlation functions and spectral densities.** Furthermore, the Wigner rotation matrix elements are related to the well-known spherical harmonics by the following expression (derived from Eqs. 4.21, 4.30 and 4.31 from (7)):

$$D^{(l)}_{0m}\,(\chi,-\theta,-\phi)=\sqrt{\frac{4\pi}{2l+1}}\;Y_{lm}(\theta,\phi)$$

,(S4)

in which *l* is the rank ($l=2$ in the present case) and $m$ is the projection of the angular momentum onto the axis of quantization. Note that $(\theta,\phi)=(\beta_{\mathrm{LP}},\alpha_{\mathrm{LP}})=(-\beta_{\mathrm{PL}},-\gamma_{\mathrm{PL}})$ where the subscripts of the Euler angles denote forward rotation from the lab to the PAS frame (LP), or inverse rotation from the PAS to the lab frame, thus the negative angles (PL) (see Fig. S1). Accordingly, the correlation functions $G_m(t)$ of the second-rank Wigner rotation matrix elements are related to the spherical-harmonic correlation functions $\tilde{G}_m(t)$ for $l=2$ by the relation: $G_m(t)=(4\pi/5)\,\tilde{G}_m(t)$, where the tilde on the right is to be noted. Here, the spherical harmonic correlation functions read:

$$G_m(t)=\left(\frac{4\pi}{2l+1}\right)\tilde{G}_m(t)$$

$$=\left(\frac{4\pi}{2l+1}\right)\langle[Y_{lm}(\theta,\phi;t)-\langle Y_{lm}(\theta,\phi)\rangle]^*\;[Y_{lm}(\theta,\phi;0)-\langle Y_{lm}(\theta,\phi)\rangle]\rangle.$$

(S5)

The spherical harmonic correlation functions $\tilde{G}_m(t)$ are often used in the literature (8-10) as an alternative to the Wigner rotation matrix correlation functions (4).

The two-sided spectral densities of the Wigner rotation matrix elements (Eq. S2) are then related to the one-sided spherical-harmonic spectral densities $\tilde{J}_m(\omega)$ with limits $(0,\infty)$ by: $J_m(\omega)=(8\pi/5)\,\tilde{J}_m(\omega)$, where the tilde on the right should again be noted. The spin-lattice relaxation rate formulated in terms of the spherical-harmonic spectral densities then becomes:

$$R_{1\mathrm{Z}}=\frac{3}{10}\pi\,\tilde{\chi}_{\mathrm{Q}}^2\left[\tilde{J}_1^{\mathrm{lab}}(\omega_0)\;+\;4\tilde{J}_2^{\mathrm{lab}}(2\omega_0)\right]$$

,(S6)

where $\tilde{\chi}_{\mathrm{Q}}=e^2qQ/\hbar=2\pi\chi_{\mathrm{Q}}$ and we have also re-introduced the "lab" superscript. Note that the results using the spherical-harmonic correlation functions and spectral densities are obtained by substituting: $(3/4)\pi^2\chi_{\mathrm{Q}}^2\;\rightarrow\;(3/10)\pi\tilde{\chi}_{\mathrm{Q}}^2$ and $J_m(\omega)\;\rightarrow\;\tilde{J}_m(\omega)$ where the tilde on the right is to be noted. Everything else is the same and it is just bookkeeping. The above formulas correspond to results found in the literature (9).

**Use of closure to describe composite motions and multiple coordinate transformations.** Next, as we have described above, it is necessary to separate the coupling Hamiltonian and correspondingly the correlation functions and spectral densities into the time-dependent and time-independent parts. The time-dependent part (nonsecular) governs the nuclear spin relaxation, while the time-independent part (secular) affects the spectral lineshape. The time dependence is expressed with respect to the director axis (the lamellar normal), while the time-independent part corresponds to the director orientation versus the laboratory axis system.

Accordingly, we can now separate the time-dependent and time-independent transformations using closure. We use closure to decompose the overall matrix elements with respect to the laboratory axes system into the time-dependent transformation with respect to the director frame, and the static or time-independent orientation of the director with respect to the laboratory frame. Use of closure readily allows us to expand any Wigner rotation matrix element into a sequence of intermediate frame transformations, or equivalently to collapse a sequence of frame transformations into the appropriate overall rotation matrix element for the overall, i.e. composite, motion.

In the present case we focus on the separation of the overall transformation from the axis system/PAS of the molecule to the laboratory frame in two steps: first the transformation to the director frame (time-dependent), and second the transformation from the director frame to the laboratory system (time-independent). Because the Wigner rotation matrix elements are members of a group, the overall rotation can be expressed in terms of other members of the group to read:

$$D_{0m}^{(2)}(\Omega_{\mathrm{PL}};t) = \sum_{p} D_{0p}^{(2)}(\Omega_{\mathrm{PD}};t)\, D_{pm}^{(2)}(\Omega_{\mathrm{DL}}) \quad .(\mathrm{S7})$$

In the above formula $\Omega_{\mathrm{PD}}(t)$ are the time-dependent Euler angles for transformation from the principal axis system (P) of the molecule to the director (D) frame, and $\Omega_{\mathrm{DL}}$ are the Euler angles for the static transformation from the director (D) to the laboratory (L) axis system.

**Director-frame spectral densities versus laboratory-frame spectral densities.** To make a comparison of the NMR relaxation times and order parameters to the results of MD simulations, we need to recognize that the time-dependent lipid fluctuations occur with respect to the frame of the director. This introduces a dependence on both the second-rank order parameter $\langle P_2 \rangle$ and the fourth-rank order parameter $\langle P_4 \rangle$ (4). Hence to compare the experimental NMR relaxation rates to the MD calculated values, we need to specify the director orientation with respect to the laboratory frame.

By applying closure (see above Eq. S7) to the overall correlation functions, and following the development through to the laboratory frame, we find that:

$$G_m^{\mathrm{lab}}(t) = \sum_{P} \left| D_{0p}^{(2)}(\Omega_{\mathrm{DL}}) \right|^2 G_p^{\mathrm{dir}}(t) \quad ,(\mathrm{S8})$$

where

$$G_p^{\mathrm{dir}}(t) = \langle [D_{0p}^{(2)}(\Omega_{\mathrm{PD}};t) - \langle D_{0p}^{(2)}(\Omega_{\mathrm{PD}})\rangle]^{*}\, [D_{0p}^{(2)}(\Omega_{\mathrm{PD}};0) - \langle D_{0p}^{(2)}(\Omega_{\mathrm{PD}})\rangle] \rangle \quad .(\mathrm{S9})$$

The director-frame correlation functions can also be written by subtracting the modulus-squared of the average value:

$$G_p^{\mathrm{dir}}(t) = \langle D_{0p}^{(2)*}(\Omega_{\mathrm{PD}}; t) D_{0p}^{(2)}(\Omega_{\mathrm{PD}}; 0)\rangle - |\langle D_{0p}^{(2)}(\Omega_{\mathrm{PD}})\rangle|^2$$

.(S10)

Similarly, for the laboratory-frame spectral densities, upon Fourier transformation we find that:

$$J_m^{\mathrm{lab}}(\omega) = \sum_P \left| D_{0p}^{(2)}(\Omega_{\mathrm{DL}}) \right|^2 J_p^{\mathrm{dir}}(\omega)$$

,(S11)

where:

$$J_p^{\mathrm{dir}}(t) = \mathrm{Re} \int_{-\infty}^{\infty} \langle [D_{0p}^{(2)}(\Omega_{\mathrm{PD}}; t) - \langle D_{0p}^{(2)}(\Omega_{\mathrm{PD}})\rangle]^* \, [D_{0p}^{(2)}(\Omega_{\mathrm{PD}}; 0) - \langle D_{0p}^{(2)}(\Omega_{\mathrm{PD}})\rangle] \, \rangle e^{-i\omega t} dt$$

.(S12)

In the above formulas the values of the Wigner rotation matrix elements $D_{0p}^{(2)}(\Omega_{\mathrm{PD}})$ can be found from the geometry of the system. The expressions for the Wigner rotation matrix elements are listed, e.g., in the Appendix of Ref. (6).

**Orientational averaging of relaxation rates.** In *randomly oriented* lipid membrane dispersions (so-called multilamellar vesicles or MLVs), the orientations where the director axis is perpendicular to the main magnetic field are most probable (because they correspond to the equator of the orientational probability distribution where the area element is maximal). For *non-oriented* (powder-type) samples, in solid-state NMR spectroscopy, the $\theta = 90°$ spectral edges (where $\theta \equiv \beta_{\mathrm{DL}}$) correspond to weak singularities (integrable) in the spectral orientational distribution function. They are the most prominent spectral features for which the relaxation rates are measured. However, because of lateral diffusion of the lipids in non-oriented powder-type distributions, i.e., multilamellar dispersions or MLVs, during the relaxation times (tens of millisecond) the relaxation rates are averaged over all director orientations. Hence, the orientation dependence of the relaxation is suppressed, or averaged (11), and one can thus assume the orientationally averaged limit.

When the director changes its orientation rapidly compared to the relaxation times, the Wigner rotation matrix elements for the director–laboratory frame transformation are averaged over the various possible values. Orientational averaging of the director with respect to the laboratory frame / *z*-axis is defined with respect to the spin-lattice relaxation times, which are typically in the range of ~50–100 msec or longer (12). In this case, the mean-square Wigner rotation matrix elements for the frame transformation are averaged to their isotropic values, leading to: $\langle |D_{0p}^{(2)}(\Omega_{\mathrm{PD}})|^2 \rangle = 1/5$. It follows that the dependence on the projection index (*m*) in the ***lab*** frame is lost due to the overall spherical symmetry. However, that does not imply the absence of a projection index (*p*) in the ***director*** frame. Quite to the contrary, the fluctuations are of limited amplitude with respect to the director as characterized by the orientational order parameters $\langle P_j(\beta_{\mathrm{PD}})\rangle$ where *j*=2, 4 in the case of NMR spectroscopy.

Hence, in the orientationally averaged case (11), the spectral densities are:

$$J_m^{\mathrm{lab}}(\omega) = \langle J_m^{\mathrm{lab}}(\omega)\rangle \equiv J(\omega) = \frac{1}{5}[J_0^{\mathrm{dir}}(\omega) + 2J_1^{\mathrm{dir}}(\omega) + 2J_2^{\mathrm{dir}}(\omega)] \tag{S13}$$

where the dependence on the projection index (*m*) in the laboratory frame vanishes due to the spherical symmetry. The spin-lattice relaxation rates are thus:

$$R_{1\mathrm{Z}} = \frac{3}{4}\pi^2\chi_{\mathrm{Q}}^2\,[J(\omega_0) + 4J(2\omega_0)] \tag{S14}$$

**Correspondence to molecular dynamics (MD) simulations.** The director-frame spectral density functions $J_p^{\mathrm{dir}}(\omega)$ describe the internal motions within the membrane and afford a direct correspondence to the results of molecular dynamics (MD) simulations. Typically, one can assume that the NMR relaxation rates are orientationally averaged as described above (for either SUVs or MLVs). The final result appropriate to molecular dynamics (MD) simulations can then be written explicitly as:

$$\begin{aligned} R_{1\mathrm{Z}} = \frac{3}{20}\pi^2\chi_{\mathrm{Q}}^2\,\{&J_0^{\mathrm{dir}}(\omega_0) + 4J_0^{\mathrm{dir}}(2\omega_0) \\ &+2[J_1^{\mathrm{dir}}(\omega_0) + 4J_1^{\mathrm{dir}}(2\omega_0)] \\ &+2[J_2^{\mathrm{dir}}(\omega_0) + 4J_2^{\mathrm{dir}}(2\omega_0)]\} \end{aligned}$$

,(S15)

where:

$$J_p^{\mathrm{dir}}(\omega) = \mathrm{Re}\int_{-\infty}^{\infty} G_p^{\mathrm{dir}}(t)e^{-i\omega t}\,dt$$

,(S16)

and:

$$G_p^{\mathrm{dir}}(t) = \langle[D_{0p}^{(2)}(\Omega_{\mathrm{PD}};t) - \langle D_{0p}^{(2)}(\Omega_{\mathrm{PD}})\rangle]^*\,[D_{0p}^{(2)}(\Omega_{\mathrm{PD}};0) - \langle D_{0p}^{(2)}(\Omega_{\mathrm{PD}})\rangle]\,\rangle$$

,(S17)

which has already been stated above. The value of the numerical pre-factor is $(3/20)\pi^2(1.70\ x\ 10^5)^2$ = $4.2785\ x\ 10^{10}\ s^{-2}$.

The above correlation function decays to a zero value because the fluctuations occur about to the average values of the Winger rotation matrix elements. The director-frame correlation functions can also be written by subtracting the modulus-squared of the average value to read:

$$G_p^{\mathrm{dir}}(t) = \langle D_{0p}^{(2)*}(\Omega_{\mathrm{PD}};t)D_{0p}^{(2)}(\Omega_{\mathrm{PD}};0)\rangle - |\langle D_{0p}^{(2)}(\Omega_{\mathrm{PD}})\rangle|^2$$

.(S18)

For a cylindrically symmetric distribution the last term on the right becomes: $|\langle D_{0p}^{(2)}(\Omega_{\mathrm{PD}})\rangle|^2\delta_{0p}$ where $\delta_{0p}$ is the Kronecker delta function. In the above correlation function the term on the left decays to a non-zero value as given by $|\langle D_{0p}^{(2)}(\Omega_{\mathrm{PD}})\rangle|^2$, which is then subtracted. These formulas lend themselves directly to molecular dynamics (MD) simulations.

**Orientationally averaged relaxation rates in terms of spherical-harmonic spectral densities.** For completeness, the results for the orientationally averaged relaxation rates can also be expressed in terms of the spherical-harmonic correlation functions (see above). Here we have the substitutions (see above): $(3/4)\pi^2\chi_{\mathrm{Q}}^2 \rightarrow (3/10)\pi\tilde{\chi}_{\mathrm{Q}}^2$ and $J_m(\omega) \rightarrow \tilde{J}_m(\omega)$ to obtain the equivalent results in terms of the spherical-harmonic correlation functions and spherical-harmonic spectral densities (indicated by the almost invisible tilde on the right that is to be noted). Everything else is the same. The result is:

$$R_{1\mathrm{Z}} = \frac{3}{50}\pi\tilde{\chi}_{\mathrm{Q}}^2 \{\tilde{J}_0^{\mathrm{dir}}(\omega_0) + 4\tilde{J}_0^{\mathrm{dir}}(2\omega_0) \\ +2[\tilde{J}_1^{\mathrm{dir}}(\omega_0) + 4\tilde{J}_1^{\mathrm{dir}}(2\omega_0)] \\ +2[\tilde{J}_2^{\mathrm{dir}}(\omega_0) + 4\tilde{J}_2^{\mathrm{dir}}(2\omega_0)]\}$$

,(S19)

[where the almost invisible tildes on the right are to be noted]. The spherical harmonic-correlation functions are given by:

$$\tilde{G}_p^{\mathrm{dir}}(t) = \langle [Y_{2p}(\Omega_{\mathrm{PD}}; t) - \langle Y_{2p}(\Omega_{\mathrm{PD}})\rangle]^* \, [Y_{2p}(\Omega_{\mathrm{PD}}; 0) - \langle Y_{2p}(\Omega_{\mathrm{PD}})\rangle] \, \rangle$$

.(S20)

In the above formula $\Omega_{\mathrm{PD}} = (0, \beta_{\mathrm{PD}}, \gamma_{\mathrm{PD}}) = (-\theta, -\phi)$ are now the angles (either Euler or spherical polar) for the transformation from PAS to director frame which are isomorphous to those in Fig. S1. The director-frame correlation functions can also be written by subtracting the modulus-squared of the average value:

$$\tilde{G}_p^{\mathrm{dir}}(t) = \langle Y_{2p}^*(\Omega_{\mathrm{PD}}; t) Y_{2p}(\Omega_{\mathrm{PD}}; 0)\rangle - |\langle Y_{2p}(\Omega_{\mathrm{PD}})\rangle|^2$$

.(S21)

where for a cylindrically symmetric distribution, the last term on the right becomes $|\langle Y_{2p}(\Omega_{\mathrm{PD}})\rangle|^2 \delta_{0p}$ where $\delta_{0p}$ is the Kronecker delta function. Note that Eqs. S20 and S21 are distinctly different from the correlation function obtained by application of the spherical harmonics addition theorem (Eq. 22 in main text) which ignores the dependence on a director axis as illustrated in Fig. 8 *A* in main text.

The results above (Eq. S15 and S19) should be used to compare MD simulations to the experimental NMR relaxation rates. In the case of MD simulations, the lipid bilayer exists as a patch. The lipid fluctuations are considered relative to the frame of the membrane patch, which we can assume is defined by the lamellar normal. The correlation functions correspond to the Wigner rotation matrix elements, which are listed in Ref. (6). To compare to the experimental NMR relaxation rates, the director frame correlation functions are used together with the orientationally averaged results for the NMR spin-lattice relaxation rate, given above.

Notably, the correlation functions $G_p^{\mathrm{dir}}(t)$ and the spectral densities $J_p^{\mathrm{dir}}(\omega)$ in closed form can be factored into their mean-square amplitudes and reduced functions/values as already indicated above. The reduced correlation functions can correspond to an exponential decay, power law, or stretched exponential decay as mentioned above, and correspondingly in the Fourier frequency domain for the reduced spectral densities. However, in the case of MD simulations the correlation functions and spectral densities are evaluated numerically. They can then be fit or tested against the simple analytical forms, both for validation and as a test of the closed-form theory.

**Reduction to the isotropic solution NMR limit.** Lastly, if one assumes unrestricted isotropic motion with a single correlation time, then the dependence on the projection index ($p$) due to the molecular principal axis vanishes because of the spherical symmetry. Summing over all the reduced spectral densities leads to the well-known isotropic solution NMR result, which reads:

$$R_{1Z} = \frac{3}{20}\pi^2\chi_Q^2\,\{j(\omega_0) + 4j(2\omega_0)\}$$

.(S22)

Here $j(\omega) = 2\tau_C/(1+\omega^2\tau_C^2)$ is a Lorentzian reduced spectral density for the molecular motions with $\tau_C$ as the single correlation time. Equivalently, the above result can be expressed using angular frequency units for the coupling constant and one-sided Lorentzian spectral densities, giving:

$$R_{1Z} = \frac{3}{40}\tilde{\chi}_Q^2\,\{\tilde{j}(\omega_0) + 4\tilde{j}(2\omega_0)\}$$

.(S23)

where $\tilde{j}(\omega) = (1/2)j(\omega) = \tau_C/(1+\omega^2\tau_C^2)$ and where the invisible tilde is again to be noted. The numerical value of the prefactor is $(3/40)(4\pi^2)(1.70\,x\,10^5)^2$ = $8.5570\,x\,10^{10}\,s^{-2}$ as stated in Ref. (9). Note that compared to solid-state NMR results (see above), the solution NMR formulas consider unrestricted rotations of the molecule, i.e., in the absence of a director axis or orientational order parameters.
